# Tumor break load quantitates structural variant-associated genomic instability with biological and clinical relevance across cancers

**DOI:** 10.1101/2024.12.08.626771

**Authors:** Soufyan Lakbir, Renske de Wit, Ino de Bruijn, Ritika Kundra, Ramyasree Madupuri, Jianjiong Gao, Nikolaus Schultz, Gerrit A. Meijer, Jaap Heringa, Remond J. A. Fijneman, Sanne Abeln

## Abstract

While structural variants (SVs) are a clear sign of genomic instability, they have not been systematically quantified per patient. Therefore, the biological and clinical impact of high numbers of SVs in patients is unknown. We introduce tumor break load (TBL), defined as the sum of unbalanced SVs, as a measure for SV-associated genomic instability. Using pan-cancer data from TCGA, PCAWG, and CCLE, we show that a high TBL is associated with significant changes in gene expression in 26/31 cancer types that consistently involve upregulation of DNA damage repair and downregulation of immune response pathways. Patients with a high TBL show a higher risk of recurrence and shorter median survival times for 5/15 cancer types. Our data demonstrate that TBL is a biologically and clinically relevant feature of genomic instability that may aid patient prognostication and treatment stratification. For the datasets analyzed in this study, TBL has been made available in cBioPortal.

## Introduction

Genomic instability is a hallmark of cancer, defined by an excessive rate of somatic DNA alterations^1^. These alterations are diverse in nature, ranging from point mutations to full genome duplications. An excessive rate of DNA alterations raises the likelihood that important genes and pathways will be affected, enabling cancer to acquire new capabilities and thereby driving tumor development^1^. Tumors exhibiting high genomic instability also demonstrate considerable inter-cellular heterogeneity, which has been associated with immune evasion and therapy resistance^2–4^. It is therefore not surprising that genomic instability has been linked to poor prognosis, metastasis, and therapeutic resistance^3,5^. At the same time, genomic instability also introduces vulnerabilities that can be exploited for treatment. For example, an excessive presentation of neo-antigens on the cell surface due to DNA mutations which the immune system can recognize makes immunotherapy more effective^6,7^. In addition, genomic instability due to deficient homologous recombination (HRD) leads to sensitivity to PARP inhibitors^8^. Therefore, measures of genomic instability have potential as both prognostic and predictive biomarkers.

Although genome instability is observed in all cancers, there is a broad spectrum in the type of instability that can occur. With the current measurement techniques, we can distinguish three main classes of genomic alterations: single/simple nucleotide variants (SNVs), somatic copy number aberrations (SCNAs), and structural variants (SVs). Most solid cancers demonstrate chromosomal instability (CIN), a form of genomic instability characterized by excessive SCNAs and SVs^3,9^. A smaller fraction of cancers show microsatellite instability (MSI), that have many SNVs^7,10^. Underlying these different types of genome instability are distinct biological mechanisms ranging from defects in DNA repair to replication stress. For example, MSI tumors have defects in DNA mismatch repair leading to the high frequency of SNVs^7^. A similar phenotype of hypermutation is displayed in cancers with *POLE or POLD1* mutations^11^. On the other hand, tumors with chromosome segregation errors can display an abundance of SCNAs^12^. Finally, defects in DNA double-strand break repair have been associated with high levels of SVs^13^. Each of these types of instabilities have been associated with different clinical outcomes that can depend on the tumor type and disease stage^5,7,14^.

To leverage the characteristics of genomic instability for clinical use, several computational quantifications have been developed. Firstly, the tumor mutational burden (TMB; **Figure 1**) quantitates the number of SNVs. In agreement with findings that link high SNV levels to greater neo-antigen presentation, the TMB has been demonstrated to be a suitable biomarker to predict response to immunotherapy^15^. Secondly, the fraction genome altered (FGA; **Figure 1**) is a measure for the extent of SCNA in the genome. Although there is no direct use of FGA in current clinical practice, this measure gives insight in the extent of aneuploidy, which has been associated with both the promotion and suppression of malignant growth^16^. SVs, however, have not yet consistently been characterized as a distinct feature of genomic instability.

**Figure 1.**
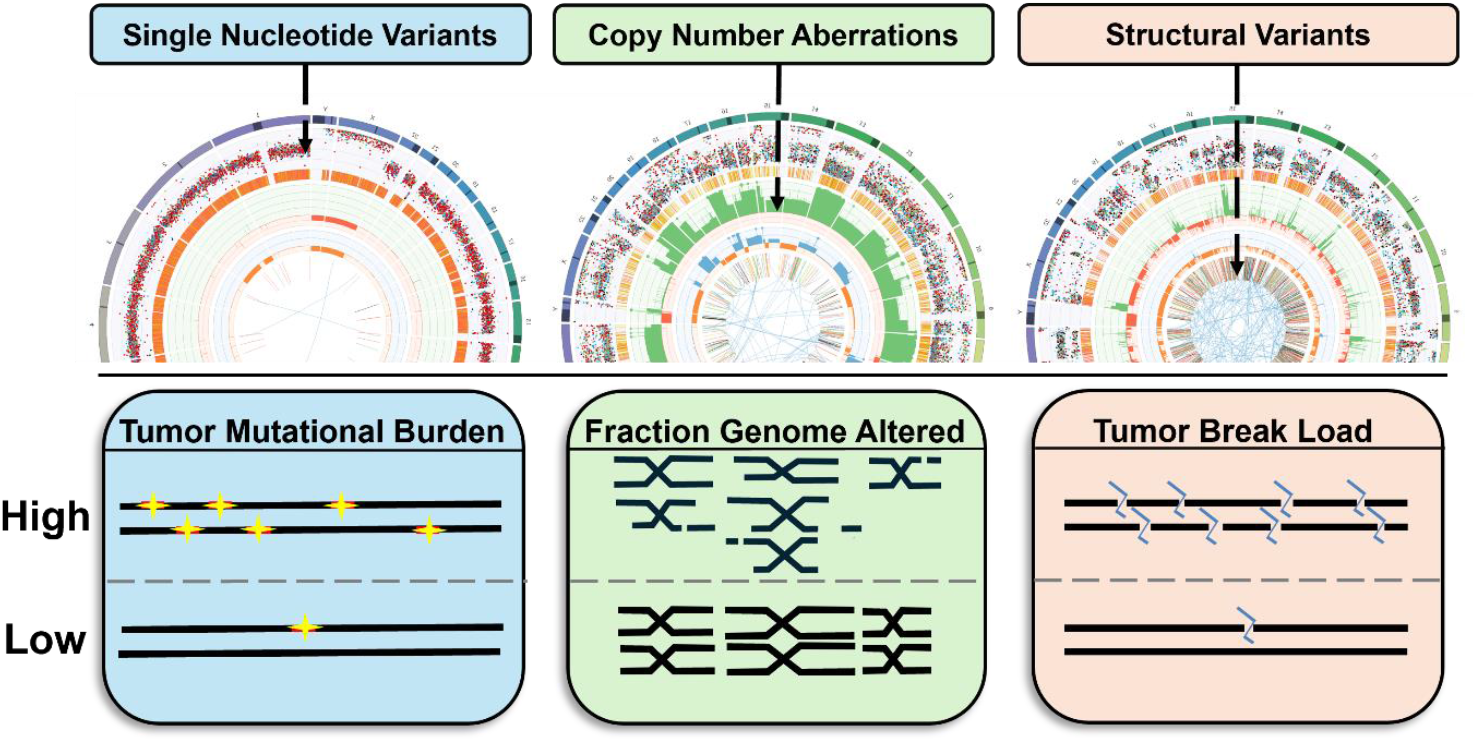
Schematic drawing of three genomic instability measures describing the extent of SNVs, i.e., the tumor mutational burden (TMB), the extent of SCNAs described by the fraction genome altered (FGA), and the extent of SVs captured in the tumor break load (TBL). Examples of CIRCOS plots from samples with high levels of each genomic instability measure are depicted above the different measures; the samples have been obtained from patients with metastatic colorectal cancer. The outermost layer shows the chromosomes. The second layer represents with each dot an SNV. The 4^th^ layer highlights DNA copy number variation with green bars representing gains and orange bars, losses. The innermost two layers show structural variants, with the bars representing breakpoints and the lines representing translocations.

Although the existence of SVs in cancer has been known for decades, they have only gained attention in recent years due to their difficulty in detection. Techniques such as comparative genome hybridization (CGH) or SNP arrays can be used to detect SVs but have insufficient resolution to detect small or balanced events^17^. Recently, the declining costs of deep whole-genome sequencing have made it feasible to perform genome-wide characterization of SVs on the nucleotide level. Although their influence on tumor biology is still insufficiently understood, SVs have already been shown to have clinical relevance. For example, tumors with homologous recombination deficiency (HRD) that harbor many SVs respond well to PARP inhibitors. However, not all tumors with many SVs show the characteristics of HRD^14^. For example, within metastatic colorectal cancer only 0.43% of tumors show the characteristics of HRD^14^. This suggests that other deficiencies can also lead to this type of genomic instability, for which the clinical consequences are still underexplored. One important reason for this has been the absence of a quantitative measure of SV burden. In previous work, we have introduced the tumor break load (TBL), a quantitative measure of the number of unbalanced SVs in the genome, as a feature of SV-associated genomic instability in colorectal cancer. We have shown that the TBL is a prognostic marker for disease-recurrence in localized microsatellite stable colorectal cancer. However, the biological and clinical implications of TBL across the full spectrum of cancer types remains to be explored.

In the present study, we aim to systematically assess the biological and clinical characteristics associated with TBL in patient and cell line data across various cancer types. We use other measures of genomic instability, TMB and FGA, to provide context for our results. Our approach is as follows (**Figure S1):** First, we characterized the landscape of genomic instability across multiple tumor types, distinguishing between three measures: TMB, FGA, and TBL. We performed this analysis in patient-derived datasets from The Cancer Genome Atlas (TCGA)^18^ and the Pan-Cancer Analysis of Whole Genomes (PCAWG)^19^, as well as cell line data from the Cancer Cell Line Encyclopedia (CCLE)^20^. Second, in the TCGA and PCAWG cohorts, we explored differences in tumor biology between samples with high and low values for each genomic instability measure. We quantified the impact on gene expression and assessed which mutations in particular genes and pathways were associated with significant differences in genomic instability measures. Finally, we investigated the relation of genomic instability measures with disease recurrence and survival.

## Results

### Wide variability in SV-associated genomic instability within and between cancer types

To gain insight into the landscape of SV-associated genomic instability across cancer, we have assessed the variability in TBL with respect to other measures of genomic instability, TMB and FGA. We made use of genomic data from primary cancer and cancer cell lines, specifically SNP6 array data from the TCGA, WGS data from the PCAWG, and WGS data from the CCLE. In the TCGA data, wide variability was observed in all three measures of genomic instability both within and between different cancer types (**Figure 2**). Overall, we observed two groups of cancer types: one with many SVs, with a median TBL above the lower quantile (14), and a group of seven cancer types with far less SVs showing a median TBL < 14. These latter cancer types also displayed a low median TMB and FGA and may not be characterized by high genomic instability in general. Uterine Corpus Endometrial Carcinoma (UCEC) showed a clear bimodal distribution for the TBL, suggesting two distinct populations with high and low SV burden. However, this bimodality was not present for the TMB and FGA. The PCAWG dataset recapitulated the wide variability observed for the TMB, FGA and TBL and also showed two groups of cancers with few and many SVs **(Figure S2)**. Although the TBL is calculated from two different data sources as described in the materials and methods, we observed high concordance in TBL (R^2^: 0.84; **Figure S3**) between shared samples from TCGA and PCAWG.

**Figure 2.**
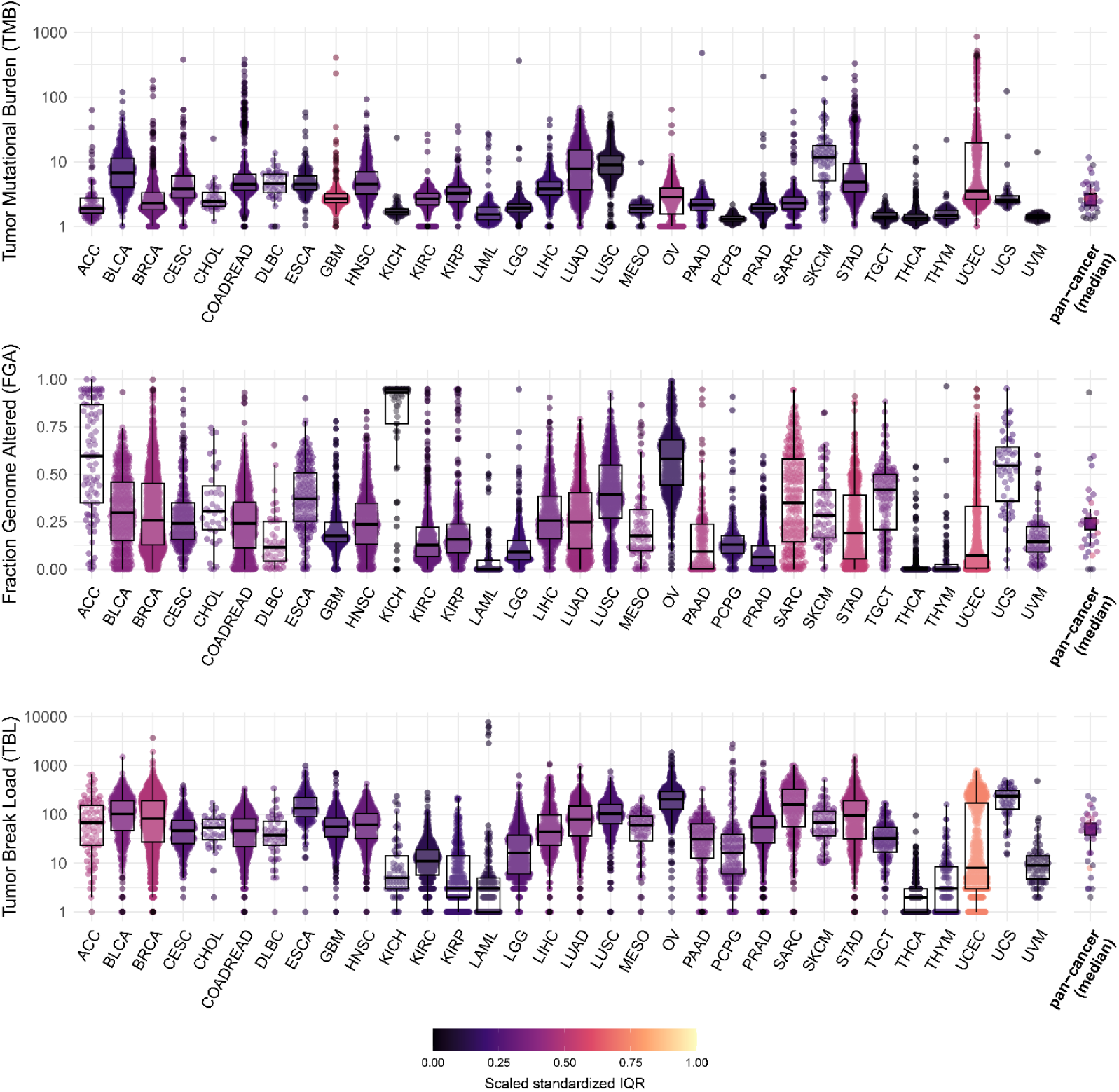
High inter- and intra-tumor type variability in genomic instability measures in cancer. Pan-cancer overview of the tumor mutational burden (TMB), the fraction genome altered (FGA) and the tumor break load (TBL) for different types of primary cancer denoted with the TCGA study abbreviations as provided in TCGA Study Abbreviations | NCI Genomic Data Commons (cancer.gov). The TMB and TBL are visualized on a log_10_ scale. The Color indicates the scaled standardized interquartile range (IQR) calculated from the ranked measures, which gives an indication of the variability in genomic instability measure per tumor type. The medians of all cancer types are reported as pan-cancer (median). ACC: Acute Myeloid Leukemia, BLCA: Bladder Urothelial Carcinoma, BRCA: Breast invasive carcinoma, CESC: Cervical squamous cell carcinoma and endocervical adenocarcinoma, CHOL: Cholangiocarcinoma, COADREAD: Colon and Rectum adenocarcinoma, DLBC: Lymphoid Neoplasm Diffuse Large B-cell Lymphoma, ESCA: Esophageal carcinoma, GBM: Glioblastoma multiforme, HNSC: Head and Neck squamous cell carcinoma, KICH: Kidney Chromophobe, KIRC: Kidney renal clear cell carcinoma, KIRP: Kidney renal papillary cell carcinoma, LAML: Acute Myeloid Leukemia, LGG: Brain Lower Grade Glioma, LIHC: Liver hepatocellular carcinoma, LUAD: Lung adenocarcinoma, LUSC: Lung squamous cell carcinoma, MESO: Mesothelioma, OV: Ovarian serous cystadenocarcinoma, PAAD: Pancreatic adenocarcinoma, PCPG: Pheochromocytoma and Paraganglioma, PRAD: Prostate adenocarcinoma, SARC: Sarcoma, SKCM: Skin Cutaneous Melanoma, STAD: Stomach adenocarcinoma, TGCT: Testicular Germ Cell Tumors, THCA: Thyroid carcinoma, THYM: Thymoma, UCEC: Uterine Corpus Endometrial Carcinoma, UCS: Uterine Carcinosarcoma, UVM: Uveal Melanoma.

To infer whether the genomic instability associated with SVs is distinct from TMB and FGA, we calculated the correlation between these three metrics using the Pearson’s correlation coefficient. Although TBL and FGA were both derived from copy number data, they showed a low correlation with each other in both primary cancer datasets and cancer cell lines **(Figure S4)**. As the FGA is a measure of the fraction of the genome copy number altered, a high FGA can occur with few breaks due to large chromosomal aneuploidies as shown in Figure S5A. In contrast, a high TBL can occur with many focal alterations resulting in a small fraction of the genome being copy number changed as shown in figure S5B. Furthermore, TBL and FGA also showed a similarly low correlation with TMB **(Figures S6 & S7)**. Hence the three genomic instability measures represent different aspects of genomic instability.

To evaluate whether the variation in TBL was correlated with clinical variables, we have assessed the association between TBL and tumor stage. TBL showed a significant correlation with tumor stage, where higher tumor stages were associated with significantly higher levels of TBL in nine out of 19 cancer types (**Figure S8**). FGA and TMB also showed a significant correlation with tumor stage in five and six tumor types, respectively (**Figure S8**). In addition, we observed that MSI tumors were associated with significantly higher TMB and lower FGA and TBL (**Figure S9**). Overall, there is wide variability in TBL between and within tumor types that shows to be a distinct feature of genomic instability, that increases along with tumor stage.

In contrast to primary cancer samples, cancer cell lines do not mimic the wide inter and intra-tumor type variation in genomic instability measures but have an overall high level of genomic instability in general compared to primary cancers (**Figure S10**). Especially the TBL measure shows high levels with a median pan-cancer TBL of 80 [69-114] breaks in the cell lines with low variation that does not match the high variability observed in patients. These results show that cancer cell lines do not recapitulate the wide variability in SV-associated genomic instability observed in primary cancers. The TBL values for all three datasets are reported in **Table S1** and are available in cBioPortal as a feature to the TCGA, PCAWG and CCLE studies.

### Distinct tumor biology characterizes high and low SV-associated genomic instability

To understand whether the wide variability in TBL observed between cancers is likewise associated with different tumor biology, we used CIBRA^21^ to quantify system-wide transcriptomic differences between tumor samples from both extremes of this genomic instability distribution. CIBRA is a computational method that leverages differential expression analysis to calculate an impact score that reflects the magnitude of variation between two conditions. For each tumor type, we defined a TBL-high and TBL-low sample group by taking the 75% quantile and the 25% quantile of the distribution (**Figure 3A**). Across cancer types, CIBRA scores revealed that high TBL was associated with significant dysregulation of the transcriptome, with 18% to 29% of genes (median 22%) being differentially expressed. This finding indicates that there are substantially different expression profiles between the TBL-high and TBL-low groups (**Figure 3B**). Subsequently, we performed the same analysis for TMB but not FGA, since FGA is a measure of the extent of SCNAs. SCNAs affect a large region of the genome, often removing or adding gene copies and affecting gene expression. As such, using gene expression to infer the impact on tumor biology of high and low FGA is biased. Like TBL, our CIBRA analysis of TMB also showed significant differences in gene expression between both extremes (median 19% [16%-22%]; **Figure 3B and Figure S11A**). However, in most cancer types (23/31), TBL displayed a higher CIBRA score than TMB, demonstrating a more pronounced distinction between the transcriptomes of the high and low TBL sample groups. Since we had observed a correlation between TBL and disease stage and MSI status (**Figure S8 and Figure S9**), we repeated the analysis for exclusively early stage (I-II) MSS samples to exclude the effect of stage and MSI, which produced highly similar results (**Figure S11B & C**). This suggests that the observed differences in gene expression are not explained by known biological variation, *i*.*e*., stage and MSI status. In the PCAWG data, the CIBRA scores were generally lower than in TCGA. Nevertheless, this dataset also showed a significant impact of high TBL on gene expression in 4/10 cancer types, such as BRCA (**Figure S12**).

**Figure 3.**
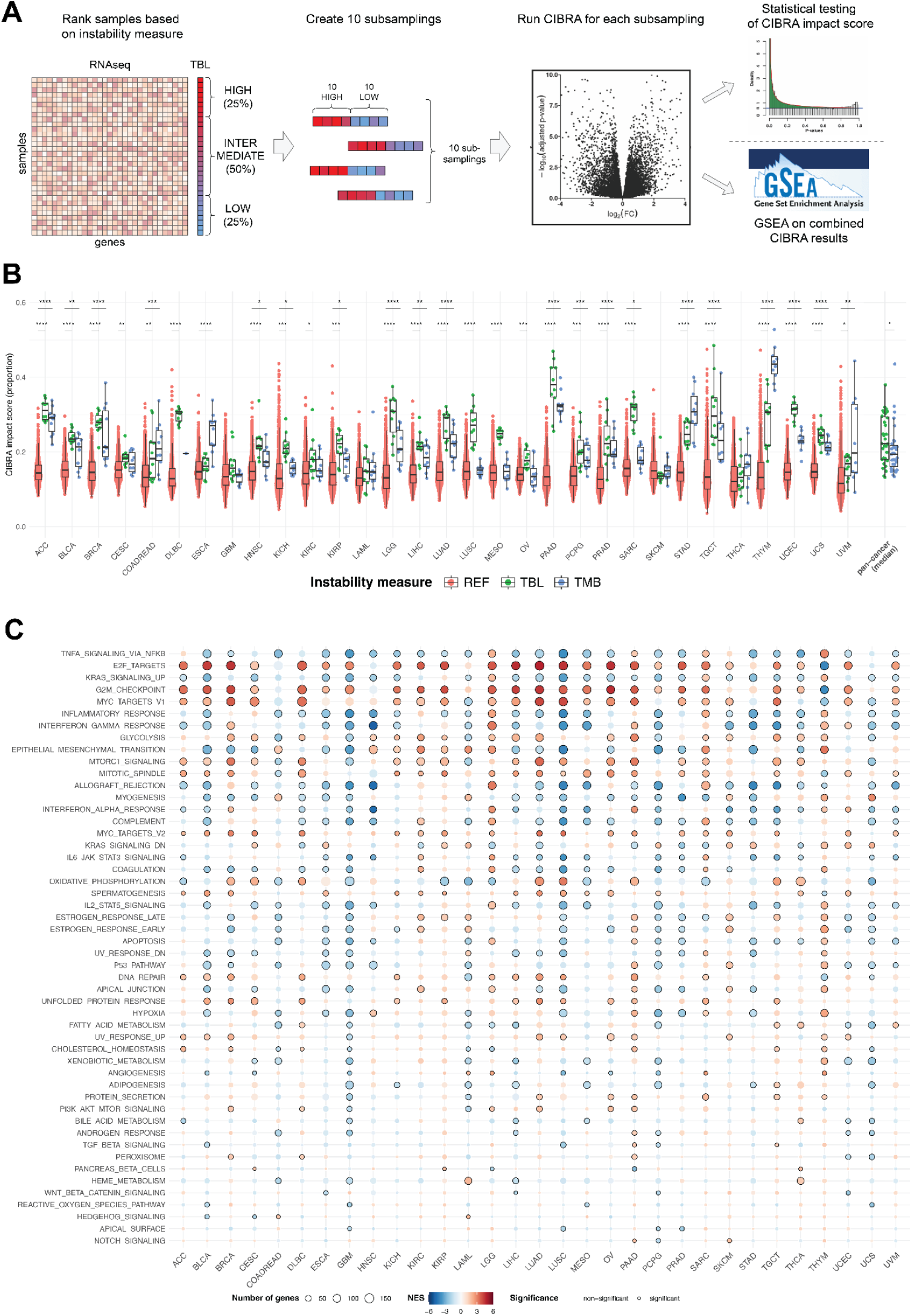
Differences in genomic instability reflect distinct tumor gene expression profiles. **A)** Overview of the comparison of gene expression between TBL-high and TBL-low tumors using CIBRA. For each tumor type, we ranked the samples from high TBL to low TBL. We identified TBL-high and TBL-low samples as the first and last quartiles of the distribution. Subsequently, we created 10 random subsamples of 10 TBL-high and 10 TBL-low samples and performed a CIBRA analysis for each subsample. The resulting CIBRA impact scores were tested for significance using a Mann-Whitney U test, compared to a reference distribution of CIBRA scores created from 1000 random permutations of the dataset. Finally, we performed gene set enrichment analysis (GSEA) on the combined CIBRA results, ranking the genes based on mean fold change and adjusted p-value across all 10 subsamples. We repeated the analysis for TMB-high and TMB-low samples. **B**) Results of the CIBRA analysis of TCGA tumor samples. Green data points indicate CIBRA scores from the TBL-high vs. TBL-low comparisons, blue points represent TMB-high vs. TMB-low, and red points depict the scores from the reference distribution, consisting of CIBRA scores from 1000 random permutations of the data separately for each cancer type. The y-axis (CIBRA proportion score) shows the fraction of genes differentially expressed between the two sample groups. In most cancer types, TBL has a higher CIBRA impact score than TMB. **C)** Results of GSEA analysis using the MSigDb Hallmark Gene Set. Pathways are ranked according to the number of cancer types in which they are differentially expressed. Colors indicate Normalized Enrichment Scores (red for up-regulated pathways, blue for down-regulated pathways). Multiple testing correction was performed using the Benjamini-Hochberg method. Black circles indicate significance (adjusted p-value < 0.01).

#### High TBL tumors are characterized by increased DNA damage repair and decreased immune response

To further interpret the differences in gene expression profiles, we performed gene set enrichment analysis (GSEA) using the MSigDb Hallmark Gene Sets, a collection of 50 gene sets reflecting cancer hallmark biological processes^22^ (**Figure 3C & Figure S14**). For TBL, we observed a consistent pattern across cancer types: high TBL was associated with significant upregulation of the E2F targets and G2M checkpoint pathways, as well as other pathways involved in cell proliferation. On the other hand, immune-related pathways (e.g. interferon gamma response) were down-regulated in TBL-high tumors. Moreover, TBL-high tumors showed an increase in the expression of DNA damage repair genes and down-regulation of the p53 pathway, which is in line with gene expression changes of high genomic instable tumors^23^. Compared to TBL, TMB showed a more heterogeneous landscape of differential gene expression, *i*.*e*. there were no pathways consistently up- or downregulated across cancer types (**Figure S13 & S15**). The upregulation of DNA damage repair pathways in TBL-high tumors was also confirmed using PCAWG data (**Figures S16 & S17**).

### Genomic alterations in DNA double strand break repair are associated with higher levels of TBL

To further gain an understanding of the distinct biological mechanisms that underly the wide variability in genomic instability measures, we assessed which genes affected by genomic alterations, *i*.*e*. SNVs, arm-level SCNAs, and SVs, were associated with significant changes in genomic instability measures across different cancer types using data from TCGA and PCAWG (**Figure 4A**). MSI tumors were excluded from this analysis, as they are a confounding factor for SNVs due to their high mutational load. SNVs affecting *TP53* were associated with significantly higher levels of TBL with an adjusted p-value < 0.05 in 12 cancer types from the TCGA and five from the PCAWG (**Figure 4; Table S2 & 3**). FGA showed a similar significant association with *TP53* mutations in the same cancer types (**Figure S18, Table S2 & 3**). However, the *TP53* mutations were associated with a significantly lower TMB in 10 cancer types (**Figure S17, Table S2 & 3**) showing that *TP53* is associated with increased chromosomal instability characterized by SCNAs and SVs, but not with an increase in the number of SNVs in the same cancer types. SVs affecting *RAD51B*, a component of the DNA double-strand break repair response, were also associated with significantly higher levels of TBL in 13 cancer types from the TCGA and in all three cancer types with an SV within *RAD51B* in the PCAWG (**Figure 4; Table S2 & 3**). In contrast, *RAD51B* SVs were associated with a significant increase in FGA in only breast (BRCA), head and neck cancer (HNSC) and lung cancer (LUAD) using data from the TCGA (**Figure S16**). Compared to *TP53* mutations, *RAD51B* SVs were more specific to a significant increase in TBL for most cancer types, highlighting its relationship with SV-associated genomic instability. While alterations affecting *TP53* and *RAD51B* were associated with increased genomic instability, alterations affecting the EGFR signaling pathway were associated with significantly lower levels of TMB (median p < 0.001) and, specifically *PIK3CA* with significantly lower levels of TBL in breast (BRCA, p < 0.00001), cervical (CESC, p < 0.01), colorectal (COADREAD, p < 0.01) and stomach (STAD, p < 0.00001) cancer (**Figure 4C, Figure S19, and Table S2 & 3**). Overall, alterations affecting genes related to DNA damage repair, specifically double strand break repair, were associated with a high TBL. Genes related to the EGFR signaling pathway were associated with low levels of genomic instability. The full catalogue of genes affected by SNVs, SVs, or arm-level SCNAs and their association with changes in TMB, FGA or TBL is shown in **supplemental table S2 and S3**.

**Figure 4.**
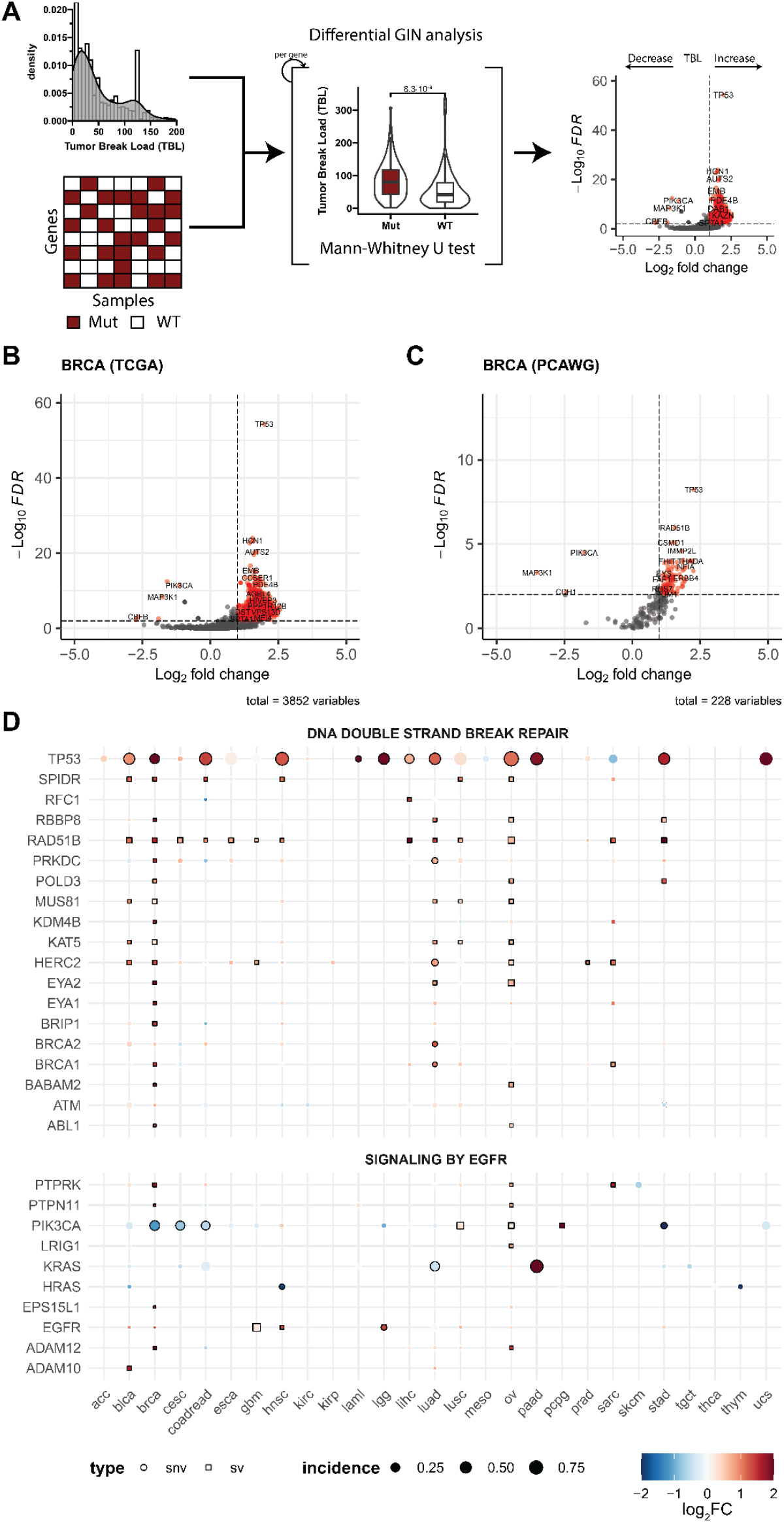
Genomic alteration association study regarding variation in TBL. **A)** Schematic overview of the workflow. A differential genomic instability measure (GIN) analysis of TBL is performed between altered and WT genes using a Mann-Whitney U test. **B-C)** Volcano plot of genes affected by SNVs, SVs, and chromosomal arms affected by SCNAs using data from primary breast cancer (BRCA) from the TCGA (**B**) and PCAWG (**C**). The x-axis represents the log_2_ foldchange in TBL between mutated and WT samples of the corresponding gene. The y-axis represents the FDR-adjusted p-value assessed with a Mann-Whitney U test. An adjusted p-value of 0.05 has been marked with a horizontal dashed line. The absolute log_2_ fold change of 1 has been marked with vertical dashed lines. **D)** Pan-cancer representation of the change in TBL with respect to genomic alterations in genes related to DNA double strand break repair and EGFR signaling using the pathway representations of Reactome. Only MSS tumors were used and only genes with a significant association with changes in the TBL in at least one cancer type are shown. Color represents the log_2_ fold change of the TBL, dot size represents the fraction of samples with a genomic alteration in the corresponding gene, and the dot shape represents the type of genomic alteration affecting the gene. The border of the dots has been colored black when the analysis was significant with an adjusted p-value < 0.05.

### High TBL is associated with a higher risk of recurrence and worse overall survival in several cancer types

Previously, we have identified TBL as a prognostic biomarker for disease recurrence in localized MSS colorectal cancer^24^. To assess whether this association is specific to colorectal cancer or a general characteristic of genomic instability, we dichotomized cancer patients into high and low TBL per tumor type using data from the TCGA. From the survival analysis of disease-free survival (DFS) using localized (stage I-III) MSS cancer data, TBL-high was associated with a significantly higher risk of disease recurrence in adrenocortical carcinoma (ACC; HR: 9.65, p = 0.0012), breast invasive carcinoma (BRCA; HR: 1.73, p = 0.03), colorectal adenocarcinoma (COADREAD; HR: 3.17, p = 0.01), kidney renal papillary cell carcinoma (KIRP; HR: 3.46, p = 0.005), and pancreatic adenocarcinoma (PAAD; HR: 6.9, p = 0.006), corrected for age, sex and tumor stage (**Figure 5, Table 1**). When we only selected patients who received either chemotherapy or radiotherapy (**Table S4**), the risk of disease recurrence became more pronounced between high and low TBL in BRCA (HR: 4.24, p = 0.01), COADREAD (HR: 10.75, p = 0.03) and PAAD (HR: 5.13, p = 0.01). A high FGA shared the higher risk of recurrence in BRCA and COADREAD and showed a significantly lower risk of recurrence in ACC and Head and Neck squamous cell carcinoma (HNSC; **Table 1**). On the other hand, high TMB was associated with a significantly lower risk of disease recurrence in BRCA and KIRP and shared the high risk of disease recurrence in ACC and COADREAD (**Table 1**). From the survival analysis using the overall survival (OS) statistic, we observed that TBL-high was associated with shorter median survival times (1113 days [642 – 1384]) in most cancers and a significantly higher risk of death (median HR 1.5 [1.1-2.4]) regardless of age, sex, and tumor stage in the majority (10/19) of cancer types (**Table S4**). Bladder urothelial carcinoma (BLCA) was the only cancer type in which a high TBL was associated with a longer median survival and a significantly lower risk of death when adjusted for age, sex, and tumor stage (0.58 HR; p = 0.001). Dichotomizing tumors in high and low FGA or TMB showed that patients with high FGA (8/19 cancer types) or TMB (7/19 cancer types) exhibited a significantly lower chance of survival (**Table S5**). However, the cancer types that showed significantly worse survival were not shared between the three measures of genomic instability. In general, TBL was a discriminatory characteristic of genomic instability in our analysis, where high values were associated with worse overall survival and a higher risk of recurrence not only in CRC but also in several other cancer types.

**Table 1.**
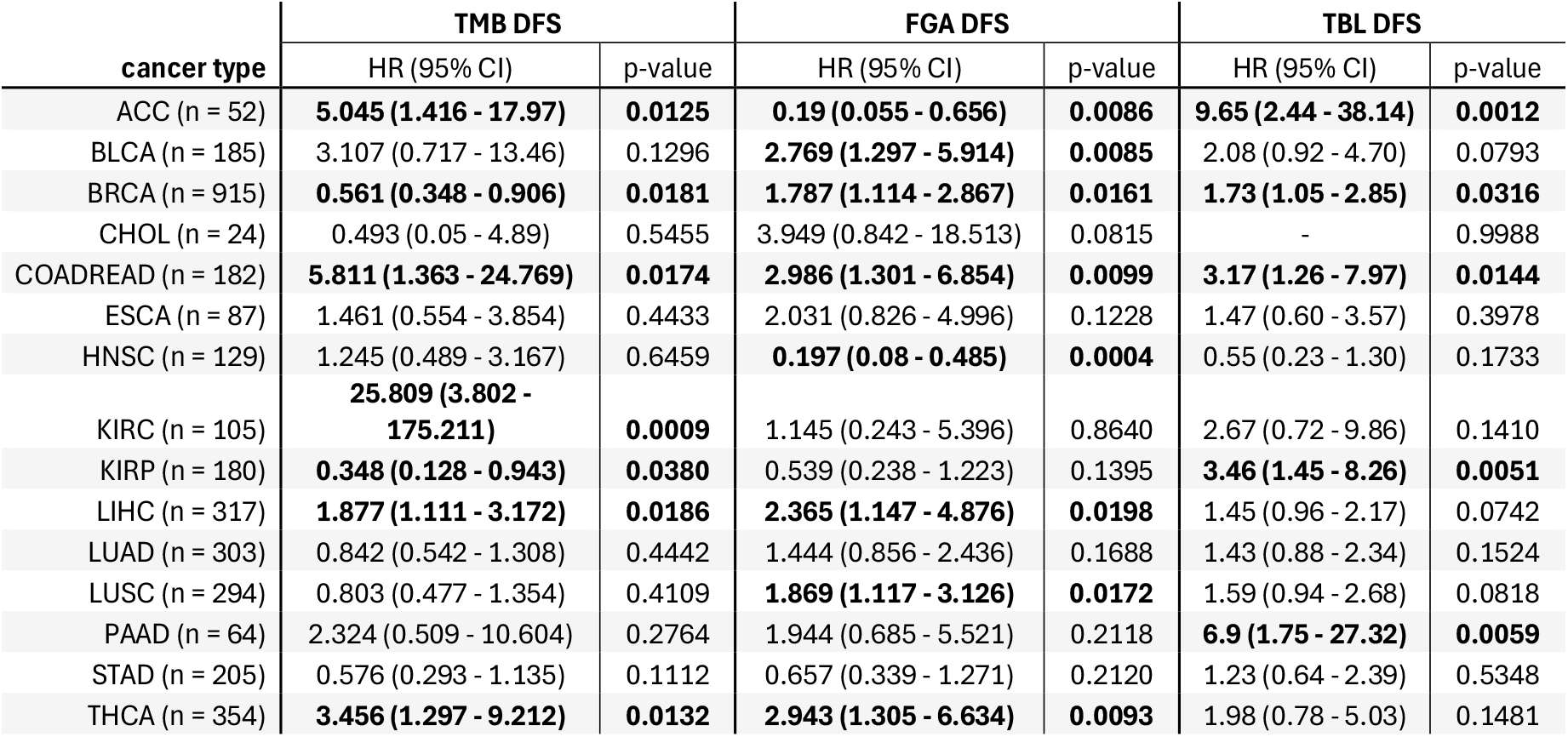
Pan-cancer survival landscape of genomic instability measures with respect to disease-free survival (DFS). Overview of the hazard rate (HR), their 95% confidence interval (95% CI) and the corresponding p-value for high compared to low TBL, FGA or TMB dichotomized using the Youden’s index as described in the methods and assessed using DFS data from localized (stage I-III) MSS cancer data from the TCGA per cancer type using a multivariate Cox proportional hazards model corrected for age, sex and tumor stages. Entries with a p-value < 0.05 are highlighted in bold. Hazard rates with an infinite confidence interval are reported as “-”. Cancer-type abbreviations are denoted with the TCGA study abbreviations as provided in TCGA Study Abbreviations | NCI Genomic Data Commons (cancer.gov).

| cancer type | TMB DFS |  | FGA DFS |  | TBL DFS |  |
| --- | --- | --- | --- | --- | --- | --- |
|  | HR (95% CI) | p-value | HR (95% CI) | p-value | HR (95% CI) | p-value |
| ACC (n = 52) | **5.045 (1.416 - 17.97)** | **0.0125** | **0.19 (0.055 - 0.656)** | **0.0086** | **9.65 (2.44 - 38.14)** | **0.0012** |
| BLCA (n = 185) | 3.107 (0.717 - 13.46) | 0.1296 | **2.769 (1.297 - 5.914)** | **0.0085** | 2.08 (0.92 - 4.70) | 0.0793 |
| BRCA (n = 915) | **0.561 (0.348 - 0.906)** | **0.0181** | **1.787 (1.114 - 2.867)** | **0.0161** | **1.73 (1.05 - 2.85)** | **0.0316** |
| CHOL (n = 24) | 0.493 (0.05 - 4.89) | 0.5455 | 3.949 (0.842 - 18.513) | 0.0815 | – | 0.9988 |
| COADREAD (n = 182) | **5.811 (1.363 - 24.769)** | **0.0174** | **2.986 (1.301 - 6.854)** | **0.0099** | **3.17 (1.26 - 7.97)** | **0.0144** |
| ESCA (n = 87) | 1.461 (0.554 - 3.854) | 0.4433 | 2.031 (0.826 - 4.996) | 0.1228 | 1.47 (0.60 - 3.57) | 0.3978 |
| HNSC (n = 129) | 1.245 (0.489 - 3.167) | 0.6459 | **0.197 (0.08 - 0.485)** | **0.0004** | 0.55 (0.23 - 1.30) | 0.1733 |
| KIRC (n = 105) | **25.809 (3.802 - 175.211)** | **0.0009** | 1.145 (0.243 - 5.396) | 0.8640 | 2.67 (0.72 - 9.86) | 0.1410 |
| KIRP (n = 180) | **0.348 (0.128 - 0.943)** | **0.0380** | 0.539 (0.238 - 1.223) | 0.1395 | **3.46 (1.45 - 8.26)** | **0.0051** |
| LIHC (n = 317) | **1.877 (1.111 - 3.172)** | **0.0186** | **2.365 (1.147 - 4.876)** | **0.0198** | 1.45 (0.96 - 2.17) | 0.0742 |
| LUAD (n = 303) | 0.842 (0.542 - 1.308) | 0.4442 | 1.444 (0.856 - 2.436) | 0.1688 | 1.43 (0.88 - 2.34) | 0.1524 |
| LUSC (n = 294) | 0.803 (0.477 - 1.354) | 0.4109 | **1.869 (1.117 - 3.126)** | **0.0172** | 1.59 (0.94 - 2.68) | 0.0818 |
| PAAD (n = 64) | 2.324 (0.509 - 10.604) | 0.2764 | 1.944 (0.685 - 5.521) | 0.2118 | **6.9 (1.75 - 27.32)** | **0.0059** |
| STAD (n = 205) | 0.576 (0.293 - 1.135) | 0.1112 | 0.657 (0.339 - 1.271) | 0.2120 | 1.23 (0.64 - 2.39) | 0.5348 |
| THCA (n = 354) | **3.456 (1.297 - 9.212)** | **0.0132** | **2.943 (1.305 - 6.634)** | **0.0093** | 1.98 (0.78 - 5.03) | 0.1481 |

**Figure 5.**
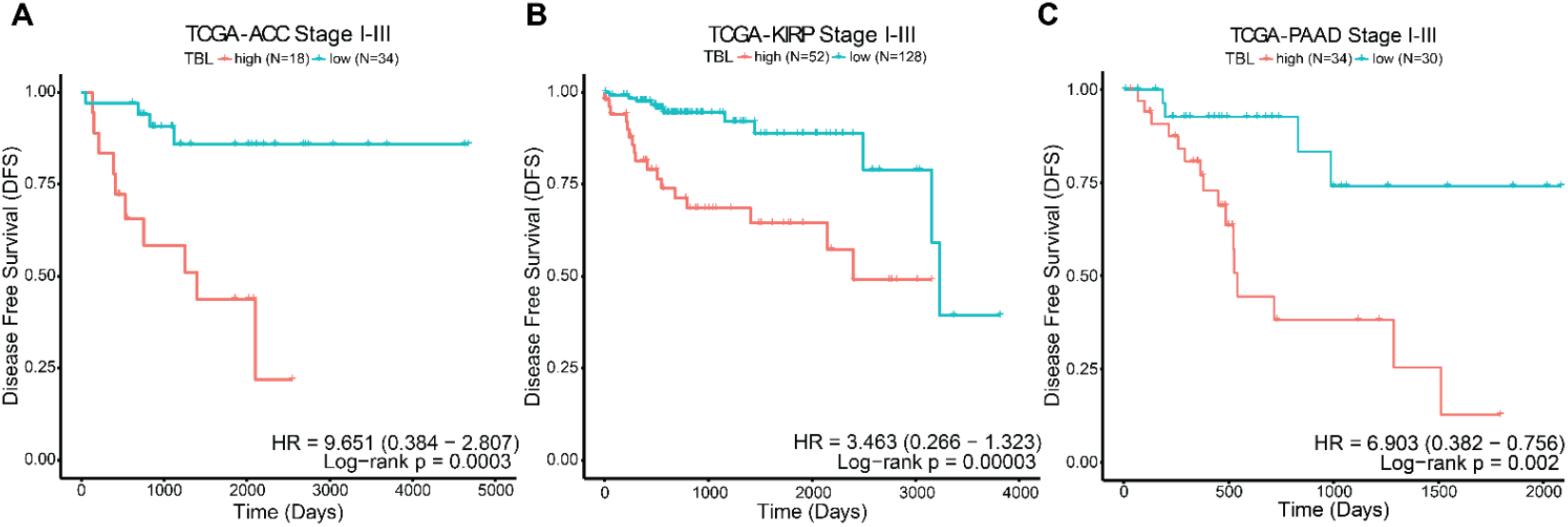
The prognostic impact of TBL in multiple cancer types. A) Kaplan–Meier Disease Free Survival (DFS) curve for 52 TCGA stage I-III Adrenocortical carcinoma (ACC) samples stratified in high and low TBL. B) Kaplan–Meier DFS curve for 180 TCGA stage I-III Kidney renal papillary cell carcinoma (KIRP) samples stratified in high and low TBL. C) Kaplan–Meier DFS curve for 64 TCGA stage I-III Pancreatic adenocarcinoma (PAAD) samples stratified in high and low TBL. Statistical significance for the difference in survival has been assessed with a log-rank test. The hazard ratio (HR) has been assessed with a multivariate Cox regression model adjusted for age, sex, and stage. TBL: Tumor Break Load; TCGA: The Cancer Genome Atlas.

## Discussion

Previously, we introduced the tumor break load (TBL) as a quantitative measure of SV burden with prognostic value for disease recurrence in the context of colorectal cancer^24^. Here, we provide a systematic assessment of the biological (*i*.*e*. associations with genomic alterations and gene expression) and clinical features of low versus high TBL across many cancer types. TBL was only weakly correlated with other measures of genomic instability, *i*.*e*. TMB and FGA, and varied widely between and within cancer types. The gene expression of samples with high TBL was significantly different from samples with low TBL across most cancer types and was characterized by high expression of DNA damage repair and decreased expression of immune-related pathways. Moreover, high TBL was associated with worse disease-free survival in many tumor types, extending our previous findings across multiple cancers. In summary, our results show that TBL captures a distinct aspect of genomic instability with prognostic value that could help patient stratification alongside other measures of genomic instability such as TMB and FGA.

Our characterization of TBL, TMB and FGA across multiple tumor types revealed large variations between and within cancer types that were generally uncorrelated between the three measures. This shows that SV-associated genomic instability is an additional, highly variable, and independent form of genomic instability in cancer. The wide variety in genomic instability measures within and between tumor types can indicate that the underlying biological mechanisms that play a role in these cancers are also highly variable. Mutational processes are influenced by many endogenous and exogenous factors, including tissue and cell type specific factors that can lead to various mutational patterns^25,26^. These patterns could be specifically exploited for prognosis or treatment, such as PARP, POLQ, or RAD51 inhibitors for targeted treatment of tumors with HRD^8^. However, individual targets are not the only way to exploit genomic instability. *General* genomic instability measures can also be applied as prognostic and predictive biomarkers, even if they represent a mixture of underlying mutational processes that lead to a similar phenotype. For example, a higher neo-antigen load or increased tumor heterogeneity are associated with better response to immunotherapy or treatment resistance, respectively^4,27^. Although both phenotypes can arise from a plethora of cellular processes, the *general* measure of genomic instability is sufficient to predict treatment response.

This study demonstrated that TBL was associated with more variation in gene expression than TMB across various cancer types. High TBL exhibited a distinct expression profile that remained consistent across most cancer types, marked by upregulation of cell cycle regulation and DNA damage repair pathways and downregulation of immune-related pathways. This suggests that the source or consequence of TBL is similar across cancer types, in contrast to the more variable tumor biology associated with extremes in TMB. The increase in alterations, characteristics of genomic instability may elicit a common response from the environment. Work from Li et al., 2023 has associated high chromosomal instability, characterized by an abundance of SCNAs and SVs, with a pro-metastatic immune microenvironment of immune-suppressive cells such as immune-suppressive macrophages^28^. Consequently, tumors with low chromosomal instability are characterized by an active immune microenvironment with CD4^+^ and CD8^+^ T-cells. In line with this work, we hypothesize that TBL-high tumors exhibit lower levels of active immune cell infiltration and have a tumor-suppressive microenvironment, which accounts for the downregulation of immune-related pathways.

In this work, we report a catalogue of genes affected by SNVs, SVs, or arm-level SCNAs and their association with changes in measures of genomic instability, which could be further explored for other new leads of genes contributing to genomic instability (**Table S2-3**). Mutations in genes involved in double-strand break repair, specifically SNVs in *TP53* and SVs in *RAD51B*, were associated with significantly higher TBL (Figure 3). Alterations in *TP53* have been reported to enable genomic instability and are associated with higher degrees of chromosomal instability, due to the loss of a critical checkpoint in the cell cycle allowing for the tolerance of accumulated damage^29,30^. Disruptions in *RAD51B*, a gene involved in homologous recombination (HR), have been associated with the accumulation of a characteristic pattern of chromosomal abnormalities that arise from HR deficiency. Furthermore, germline variants in *RAD51B* have been linked to susceptibility to breast and ovarian cancer^31,32^. These findings highlight that different genomic alterations related to DNA damage repair can lead to the same increase in TBL. However, these alterations do not fully describe the TBL, as this measure is a general feature of SV-associated genomic instability and may be influenced by multiple SV-inducing processes, like HR deficiency, chromosome segregation errors, and replication stress^33^. The impact of somatic SVs in inducing genomic instability should be further explored. In contrast to the association of *TP53* and *RAD51B* with high TBL, alterations in the EGFR pathway showed an association with low genomic instability measures. The high proliferative rate of mutant *EGFR* pathway tumors could induce a high rate of replicative stress that, in combination with high genomic instability, might push the tumor over the limit of tolerating DNA damage, resulting in *EGFR* pathway mutant tumors to predominantly display low levels of genomic instability^34,35^. This finding also highlights the potential vulnerability of *EGFR* mutant tumors in combination with DNA damage-inducing treatments^35,36^.

Genomic instability is associated with worse overall survival and a higher chance of metastasis. However it can also offer therapeutic potential, exemplified by the application of PARP inhibitors in HR-deficient tumors and the efficacy of immunotherapy in MSI cancers ^3,5,14,27^. In our study, high genomic instability measures were correlated with significantly shorter survival times. Similarly, high TBL was associated with a higher risk of recurrence in several tumor types, confirming the association of genomic instability with a worse prognosis in a variety of cancer types. Metastasized tumors have been reported to have a higher number of SVs compared to primary tumors^37^. This aligns with our observation that the TBL increases with disease stage and high TBL is associated with a higher risk of disease recurrence. This suggests that it may be a characteristic that is associated with metastatic spread. The high number of SVs may lead to the activation of the cGAS-STING pathway, facilitating migration, invasion, and metastasis^38,39^. In this way, a high TBL could be associated with an increased likelihood of developing metastases and relapse of the disease. However, within our analysis there were no clear changes in gene expression associated with cGAS-STING. Our findings reinforce the prognostic value of TBL as a biomarker for disease recurrence, extending beyond colorectal cancer to other cancer types, and show that TBL alongside FGA and TMB have potential as a general biomarker for disease recurrence across cancer types^24^. Further validation is needed to verify its prognostic value in other cancer types. As the TBL can be derived from segmented copy number profiles generated with SNP6 arrays and low-coverage WGS, or SV calls from WGS data, there is potential for wide implementation in prospective and retrospective studies.

In conclusion, our data demonstrate that TBL is a biologically and clinically relevant feature of genomic instability that may aid patient prognostication and treatment stratification in addition to the FGA and TMB, pending further validation.

## Materials & Methods

### Datasets and genomic instability metrics

To assess the clinical and biological implications of TBL in relation to other measures of genomic instability across various cancers, public data from two primary pan-cancer datasets with genomics, transcriptomics, and clinical data were used, i.e. The Cancer Genome Atlas (TCGA) and the Pan-Cancer Analysis of Whole Genomes (PCAWG). To infer how patient characteristics relate to genomic instability characteristics in cancer cell lines, data from the Cancer Cell Line Encyclopedia (CCLE) were used.

TCGA data were collected from the Genome Data Commons (GDC) portal for 33 cancer types^18^. Available processed single nucleotide variant calls using whole-exome sequencing data were retrieved using the R package TCGAbiolinks (version 2.25.3, RRID:SCR_017683)^40^, with query parameters: data.category = “Simple Nucleotide Variation”, data.type = “Masked Somatic Mutation”, legacy = FALSE, access = “open” and workflow.type = “Aliquot Ensemble Somatic Variant Merging and Masking”. Available RNA-Seq data were retrieved with the query parameters: data.category = “Transcriptome Profiling”, data.type = “Gene Expression Quantification” and workflow.type = “STAR – Counts”. Identifier mapping was performed using the R package TCGAutils (version 1.22.2). Only unique identifiers were preserved. Clinical data, the tumor mutational burden (TMB), and the fraction genome altered (FGA) were retrieved from cBioPortal^41–43^. ‘Silent’ variants indicated by the variant classification provided in the Mutation Annotation Format (MAF) file were removed from the genome-wide screen analysis. Copy-number associated SVs were called using the processed SNP6 data as described in Lakbir et al 2022^24^. In short, chromosomal breaks were defined as the boundary between two adjacent segments using ‘masked’ DNA copy number segmented data. Two parameters were used to select likely somatic chromosomal breaks: the smallest adjacent segment size (SAS) and the break size (BrS) with the thresholds SAS < 20 probes and BrS < 0.135 copy number. Any chromosomal break detected in normal samples was removed from the filtered set of chromosomal breaks. The tumor break load (TBL) was defined as the sum of SCNA-associated SVs, *i*.*e*., the sum of unbalanced somatic chromosomal breaks per tumor sample.

Public data from PCAWG were retrieved from the ICGC Data Portal (https://dcc.icgc.org/releases/PCAWG)^19^. Consensus structural variant (SV) and copy number variation (CNV) data provided by PCAWG were used for SV and SCNA calls. The SNV calls provided for the ICGC patients of the PCAWG were retrieved from the ICGC Data Portal and appended with the SNV calls from the TCGA patients retrieved from the GDC portal. The TMB and FGA were retrieved from cBioPortal (https://www.cbioportal.org/study/summary?id=pancan_pcawg_2020). The TBL was calculated by summing the number of unbalanced SVs, *i*.*e*. the deletions and duplications, per tumor sample. Only SVs that passed all filter criteria for the consensus pipeline were considered.

Public data from the CCLE were retrieved from cBioPortal (https://www.cbioportal.org/study/summary?id=ccle_broad_2019) and SV data from the depmap portal^20^. The TMB and FGA were retrieved from cBioPortal. The TBL was calculated by summing the number of unbalanced SVs, *i*.*e*. the deletions and duplications, per tumor sample.

### Inter and intra tumor variability in genomic instability

To assess the inter and intra tumor variability, *i*.*e*. the variation in genomic instability measures across and within different cancer types, we first ranked the measures in the entire pan-cancer dataset per genomic instability measure. On those ranks, we calculated the interquartile range per tumor type and normalized those values by dividing by the total range of ranks. In this way, we calculated a standardized interquartile range that can be compared between genomic instability measures with different ranges.

### Association analysis between genomic instability measures, stage, and microsatellite instability status

To understand the extent of the information shared between the different measures of genomic instability, the Pearson’s correlation coefficient (R^2^) was calculated using the ‘cor’ function from R. Additionally, the association of different measures of genomic instability with stage and microsatellite instability status was statistically evaluated using data from the TCGA. Stage as reported under “Genomic Commons Neoplasia Stage” in the clinical data was collapsed into Stage I, II, III, and IV. Per cancer type, a Mann-Whitney U test was used to assess differences in TMB, TBL, and FGA between the 4 stages, as well as between MSS and MSI samples.

### Pan-cancer genomic instability CIBRA analysis

To assess whether differences in genomic instability measures also reflect system-wide changes in gene expression, a pan-cancer CIBRA screen was performed. For TCGA, only samples were selected corresponding to primary tumors for which RNA-Seq data was available.

Per cancer type and genomic instability measure, samples were ranked in descending order and subsequently annotated as “HIGH” (first quartile), “LOW” (last quartile), or “INTERMEDIATE” (second and third quartiles) for that instability measure. Only cancer types for which at least 10 “HIGH” samples were available were used for CIBRA analysis.

CIBRA impact scores are dependent on the number of samples that were analyzed. To enable comparison of scores between different cancer types, while still preserving the variation in transcriptomic profiles within one cancer type, CIBRA analysis was performed multiple times on random subsets of 10 samples. For each combination of cancer type and instability measure, 10 random subsamples of 10 “HIGH” and 10 “LOW” samples were created. If the cancer type only had 10 “HIGH” and “LOW” samples available, only one CIBRA run was performed, as this dataset did not allow for the creation of multiple non-identical subsets. Note that partial overlap was possible between the different subsamples.

After the creation of the subsample datasets, CIBRA analysis was performed for each of the subsamples using a custom, in-house version of CIBRA adapted from CIBRA version 1.0 (Lakbir et al., 2024)^44^. The only modifications compared to the published script were the inclusion of logging statements for debugging. DESeq2 (version 1.40.2; RRID: SCR_015687)^45^ was used for the differential expression analysis that underlies the CIBRA impact score.

Statistical evaluation of the CIBRA impact scores was performed by comparing the scores of the different subsamples with a reference distribution of 1000 CIBRA runs corresponding to random permutations of the dataset for that cancer type (10 cases, 10 controls). Significance was assessed using the Mann-Whitney U test.

CIBRA analysis for the PCAWG dataset was performed in the same manner, except that, due to the low number of samples, only 100 permutations per cancer type were performed for the reference distribution.

### Gene set enrichment analysis of CIBRA results

To interpret the differences in gene expression underlying the CIBRA impact scores, gene set enrichment analysis (GSEA) was performed on the CIBRA results averaged over all subsamples. For each cancer type and genomic instability measure, the genes were ranked according to a score calculated according to the following formula: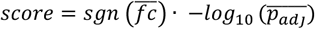 Where 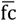 is the mean fold change across all subsamples and 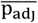 is the mean adjusted p value across all subsamples.

Hence, the genes were ordered from upregulated with highest significance, to downregulated with highest significance, with non-differentially expressed genes in the center.

Subsequently, GSEA was performed using the R package clusterProfiler (version 4.10.1; RRID:SCR_016884)^46^. Genes were annotated with the MSigDB Hallmark Gene Set^22^, using its implementation in the R package msigdbr (version 7.5.1; RRID:SCR_022870)^47^.

### Association analysis between genomic alterations and the extent of genomic instability

To identify which genomic alterations are associated with a change in genomic instability, a differential analysis was performed. The analysis was performed per gene and type of alteration (SNVs, SCNAs, and SVs). Only coding SNVs were considered in the analysis. For SVs, using the PCAWG calls, the breakpoint position was taken to match the breakpoints calculated from the TCGA SNP6 data. To reduce the number of highly correlated events, SCNAs were not assessed per gene but evaluated on chromosome arm level as provided by cBioPortal for the TCGA and GISTIC for the PCAWG dataset. SCNAs were separately assessed for gains (log_2_ copy number ratio > 0.2) and losses (log_2_ copy number ratio < -0.2). Genomic alterations were mapped to genes using AnnotationHub (version 3.8.0, RRID:SCR_024227) with queries “AH10684” and “AH110869” for PCAWG and TCGA data respectively. To reduce the number of false positive hits, MSI tumors were removed from the analysis, since they are characterized by an excessive number of SNVs.

Differences in TMB, FGA, and TBL were statistically assessed using a Mann-Whitney U test. This nonparametric test was chosen since all three genomic instability measures are not normally distributed. Moreover, only genes with a minimum of 10 mutated samples and 10 non-mutated samples were considered, given the power of the Mann-Whitney U test. The sample size needed given an alpha of 0.05 was calculated using the following example. If we consider 20000 coding genes affected by an alteration as possible tests (m), and a Bonferroni multiple testing correction, a minimal significant p-value needed given an α of 0.05 is 2.5e-06, *i*.*e*., 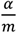.^48^ To determine the minimal sample size needed for this power, we made use of the sample size calculation described in Zhao et al. (2008)^49^ and determined a minimum of 10 samples in both groups were needed.

To account for biases due to the local mutational load within the gene of interest and its association with measures that describe the genome-wide mutational load, e.g. TMB for SNVs, a correction was made. For the association between SNVs and the TMB, since the TMB represents the extent of genome wide SNVs, the number of SNVs within the gene of interest in a sample was subtracted from the TMB for that sample. A similar correction was performed for SVs and the TBL.

### Survival analysis

To assess the clinical impact of TBL, TMB, and FGA, a threshold was determined to divide tumor samples from clinical series into ‘high’ and ‘low’ for the different measures. The Youden’s index determined from ROC-curves with the coords function (pROC package, version 1.18.5, RRID:SCR_024286)^50^ has been used as threshold. Kaplan-Meier analysis and Cox proportional-hazards regression models were applied to assess the association of the pre-defined dichotomized states with overall survival (OS) and disease-free survival (DFS) data from the TCGA using the R package survminer (version 0.4.9, with default parameters; RRID:SCR_021094). For DFS data, only localized (stage I-III) untreated samples were considered. Treatment information was retrieved using the TCGAbiolinks R package (version 2.28.4, RRID:SCR_017683) with the GDCquery function with parameters: data.category = “clinical” and data.type = “Clinical Supplement” and the GDCprepare_clinic function with parameter clinical.info = “drug”. A multivariate Cox proportional-hazards regression model was built to assess the association between the dichotomized states and age, sex, and tumor stage for data from the TCGA. Log-rank test p-values and hazard ratios (HR) were calculated using the R package survival (version 3.6-4, with default parameters; RRID:SCR_021137)^51^. A p-value < 0.05 was considered significant.

## Data availability

All data generated and analyzed during this study are publicly available and included in this published article and its supplementary files. The TBL data is also available in the public TCGA, PCAWG and CCLE studies of cBioPortal (https://cbioportal.org).

## Code availability

The underlying code for this study is available at https://github.com/nayfous/TBL_pancan_paper

## Acknowledgements

This study was supported by PPP Allowance [LSHM19027 and LSHM21018] made available by Health∼Holland, Top Sector Life Sciences & Health, to stimulate public-private partnerships. The funder played no role in study design, data collection, analysis and interpretation of the data, or the writing of this manuscript. The authors would like to acknowledge the Research High Performance Computing (RHPC) facility of the Netherlands Cancer Institute (NKI).

We thank Steven Ketelaars and Carmen Rubio-Alarcón for their valuable input on the interpretation of the immune pathway results and survival data analysis.

## Author contributions

S.L. has contributed to the conceptualization, formal analysis, validation, investigation, visualization, methodology, writing of the original draft. R.W. has contributed to the conceptualization, formal analysis, validation, investigation, visualization, methodology, writing of the original draft. I.B. has contributed to data curation and review of the writing. R.K. has contributed to data curation and review of the writing. R.M. has contributed to data curation and review of the writing. J.G. has contributed to data curation and review of the writing. N.S. has contributed to data curation and review of the writing. G.A.M. and J.H. have contributed to the conceptualization, supervision, review of the writing, and funding acquisition. R.J.A.F. has contributed to the conceptualization, supervision, funding acquisition, writing of the original draft, and review of the writing. S.A. has contributed to the conceptualization, supervision, funding acquisition, writing of the original draft, review of the writing, and methodology.

