## Supplementary material for "Tumor break load quantitates structural variant-associated genomic instability with biological and clinical relevance across cancers": supplementary_data.pdf

### Supplemental figures

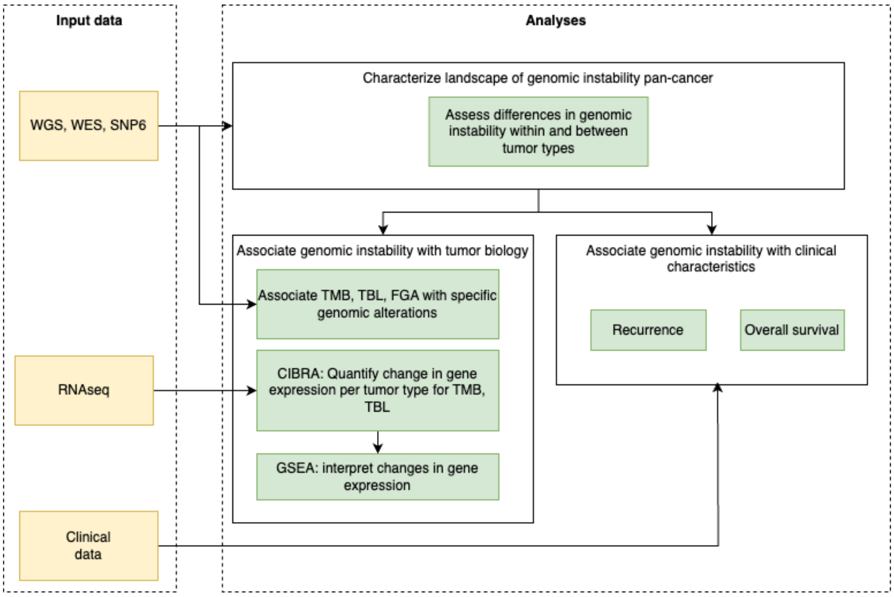


**Figure S1** Overview of the workflow used in this study with data from the TCGA, PCAWG and CCLE. First, we determined the value of different genomic instability measures (TMB, FGA, TBL) per sample, and characterized the landscape of genomic instability across different tumor types. Second, we evaluated the association of TBL (compared to TMB) with tumor biology, namely with specific genomic alterations and changes in gene expression (CIBRA, GSEA). Finally, we related differences in genomic instability measures to clinical characteristics.

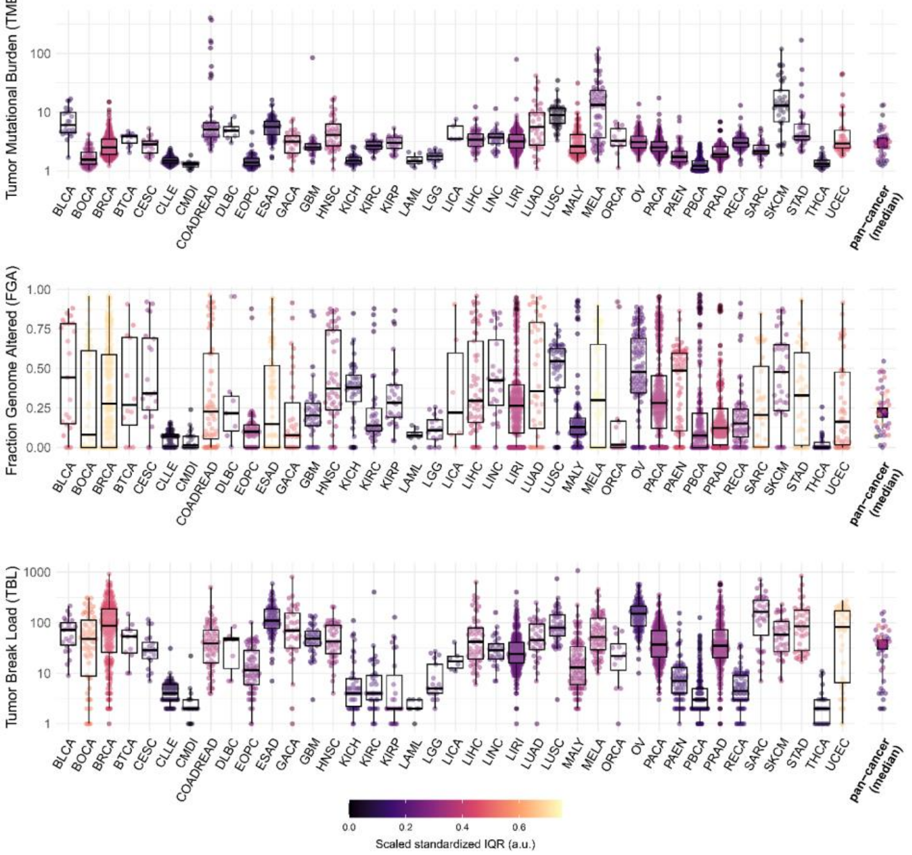


**Figure S2**. **High inter and intra variability in genomic instability measures in cancer.** Pan-cancer overview of tumor mutational burden (TMB), fraction genome altered (FGA), and tumor break load (TBL) in primary cancer (PCAWG). TMB and TBL are visualized on a $\log_{10}$ scale. The color indicates the scaled standardized interquartile range (IQR) calculated from the ranked measures. The medians of all cancer types are reported as pan-cancer (median). BLCA: Bladder Urothelial Carcinoma, BOCA: Bone Cancer, BRCA: Breast invasive carcinoma, BTCA: Biliary tract cancer, CESC: Cervical squamous cell carcinoma and endocervical adenocarcinoma, CLLE: Chronic Lymphocytic Leukemia, CMDI: Chronic Myeloid Disorders, COADREAD: Colon and Rectum adenocarcinoma, DLBC: Lymphoid Neoplasm Diffuse Large B-cell Lymphoma, EOPC: Early Onset Prostate Cancer, ESAD: Esophageal Adenocarcinoma, GACA: Gastric Cancer, GBM: Glioblastoma multiforme, HNSC: Head and Neck squamous cell carcinoma, KICH: Kidney Chromophobe, KIRC: Kidney renal clear cell carcinoma, KIRP: Kidney renal papillary cell carcinoma, LAML: Acute Myeloid Leukemia, LGG: Brain Lower Grade Glioma, LICA: Liver Cancer, LIHC: Liver hepatocellular carcinoma, LINC: Liver Cancer, LIRI: Liver Cancer, LUAD: Lung adenocarcinoma, LUSC: Lung squamous cell carcinoma, MALY: Malignant Lymphoma, MELA: Melanoma, ORCA: Oral Cancer OV: Ovarian serous cystadenocarcinoma, PACA: Pancreatic Cancer, PAEN: Pancreatic Cancer Endocrine neoplasms, PBCA: Pediatric Brain Cancer, PRAD: Prostate adenocarcinoma, RECA: Renal Cancer, SARC: Sarcoma, SKCM: Skin Cutaneous Melanoma, STAD: Stomach adenocarcinoma, THCA: Thyroid carcinoma, UCEC: Uterine Corpus Endometrial Carcinoma.

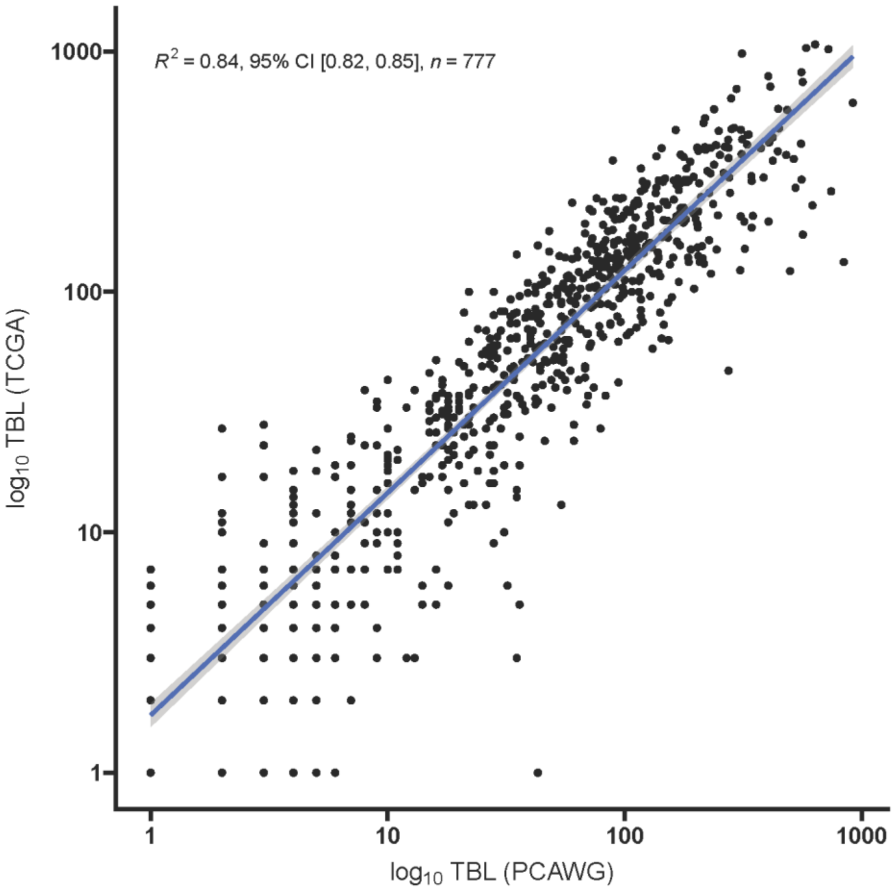


**Figure S3**. **High concordance between SNP6 data derived TBL and WGS data derived TBL.** Scatterplot of the $\log_{10}$ TBL calculated for shared samples from SNP6 array data from the TCGA and WGS data from the PCWAG. Regression line with the Pearsons' correlation coefficient ($R^2$) reported with its 95% confidence interval (CI).

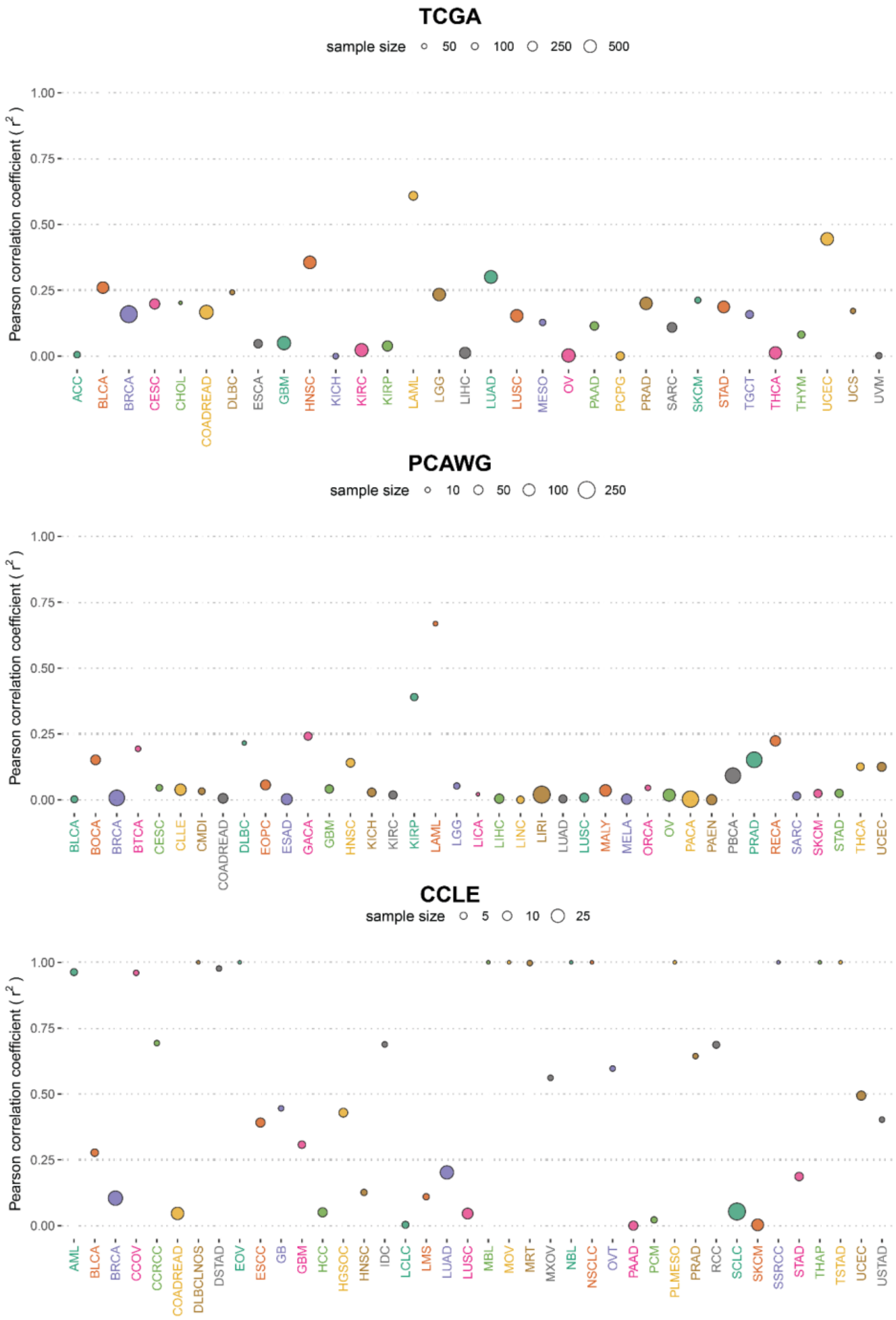
TCGA
sample size 50 100 250 500
1.00
0.75
0.50
0.25
0.00
Pearson correlation coefficient ( r² )
ACC BLCA BRCA CESC CHOL COADREAD DLBC ESCA GBM HNSC KICH KIRC KIRP LAML LGG LIHC LUAD LUSC MESO OV PAAD PCPG PRAD SARC SKCM STAD TGCT THCA THYM UCEC UCS UVM
PCAWG
sample size 10 50 100 250
1.00
0.75
0.50
0.25
0.00
Pearson correlation coefficient ( r² )
BLCA BOCA BRCA BTCA CESC CLLE CMDI COADREAD DLBC EOPC ESAD GACA GBM HNSC KICH KIRC KIRP LAML LGG LICA LIHC LINC LIRI LUAD LUSC MALY MELA ORCA OV PACA PAEN PBCA PRAD RECA SARC SKCM STAD THCA UCEC
CCLE
sample size 5 10 25
1.00
0.75
0.50
0.25
0.00
Pearson correlation coefficient ( r² )
AML BLCA BRCA CCOV CCRCC COADREAD DLBCLNOS DSTAD EOV ESCC GB GBM HCC HGSOC HNSC IDC LCLC LMS LUAD LUSC MBL MOV MRT MXOV NBL NSCLC OVT PAAD PCM PLMESO PRAD RCC SCLC SKCM SSRCC STAD THAP TSTAD UCEC USTAD

**Figure S4. TBL and FGA are distinct features of genomic instability in primary cancer.** Pan-cancer overview of the correlation between TBL and FGA reported with Pearson's correlation coefficient ($r^2$) for primary cancer data (PCAWG and TCGA) and cell lines (CCLE). The size of the dot indicates the sample size. Color marks the type of cancer. Tumor type abbreviations from the CCLE cell lines were retrieved from OncoTree. ACC: Acute Myeloid Leukemia, BLCA: Bladder Urothelial Carcinoma, BRCA: Breast invasive carcinoma, CESC: Cervical squamous cell carcinoma and endocervical adenocarcinoma, CHOL: Cholangiocarcinoma, COADREAD: Colon and Rectum adenocarcinoma, DLBC: Lymphoid Neoplasm Diffuse Large B-cell Lymphoma , ESCA: Esophageal carcinoma, GBM: Glioblastoma multiforme, HNSC: Head and Neck squamous cell carcinoma, KICH: Kidney Chromophobe , KIRC: Kidney renal clear cell carcinoma, KIRP: Kidney renal papillary cell carcinoma, LAML: Acute Myeloid Leukemia, LGG: Brain Lower Grade Glioma, LIHC: Liver hepatocellular carcinoma, LUAD: Lung adenocarcinoma, LUSC: Lung squamous cell carcinoma, MESO: Mesothelioma , OV: Ovarian serous cystadenocarcinoma, PAAD: Pancreatic adenocarcinoma, PCPG: Pheochromocytoma and Paraganglioma, PRAD: Prostate adenocarcinoma, SARC: Sarcoma, SKCM: Skin Cutaneous Melanoma, STAD: Stomach adenocarcinoma, TGCT: Testicular Germ Cell Tumors, THCA: Thyroid carcinoma, THYM: Thymoma, UCEC: Uterine Corpus Endometrial Carcinoma, UCS: Uterine Carcinosarcoma, UVM: Uveal Melanoma. BLCA: Bladder Urothelial Carcinoma, BOCA: Bone Cancer, BTCA: Biliary tract cancer, CLLE: Chronic Lymphocytic Leukemia, CMDI: Chronic Myeloid Disorders, DLBC: Lymphoid Neoplasm Diffuse Large B-cell Lymphoma, EOPC: Early Onset Prostate Cancer, ESAD: Esophageal Adenocarcinoma, GACA: Gastric Cancer, KICH: Kidney Chromophobe, LICA: Liver Cancer, LINC: Liver Cancer, LIRI: Liver Cancer, MALY: Malignant Lymphoma, MELA: Melanoma, ORCA: Oral Cancer  OV: Ovarian serous cystadenocarcinoma, PACA: Pancreatic Cancer, PAEN: Pancreatic Cancer Endocrine neoplasms, PBCA: Pediatric Brain Cancer, RECA: Renal Cancer, UCEC: Uterine Corpus Endometrial Carcinoma.

**Commented [RF1]:** Headers: TCGA – PCAWG – CCLE (instead of 'cell lines')… be consistent!!

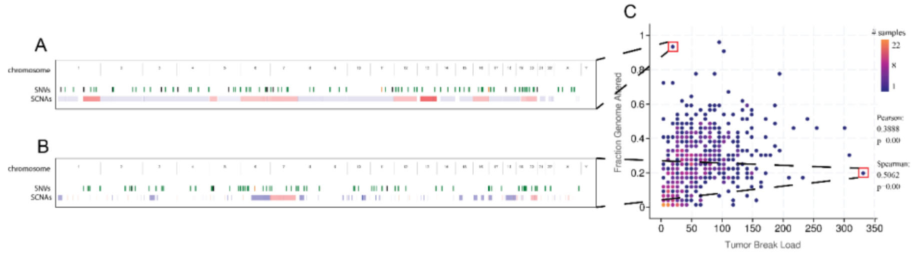


**Figure S5**. cBioPortal visualization of molecular profiles of patients with a high fraction genome altered (FGA) and low tumor break load (TBL) (A) or low FGA and high TBL (B) selected from the study view of TCGA-COADREAD depicting the tumor break load against the fraction genome altered (C). The molecular profile shows a genome-wide view of the single nucleotide variants (SNV) in green and the somatic copy number aberrations (SCNA) in blue (loss) or red (gain). The color in the scatterplot indicates the sample density. The correlation between the TBL and FGA has been reported with the Pearson and Spearman correlation coefficients.

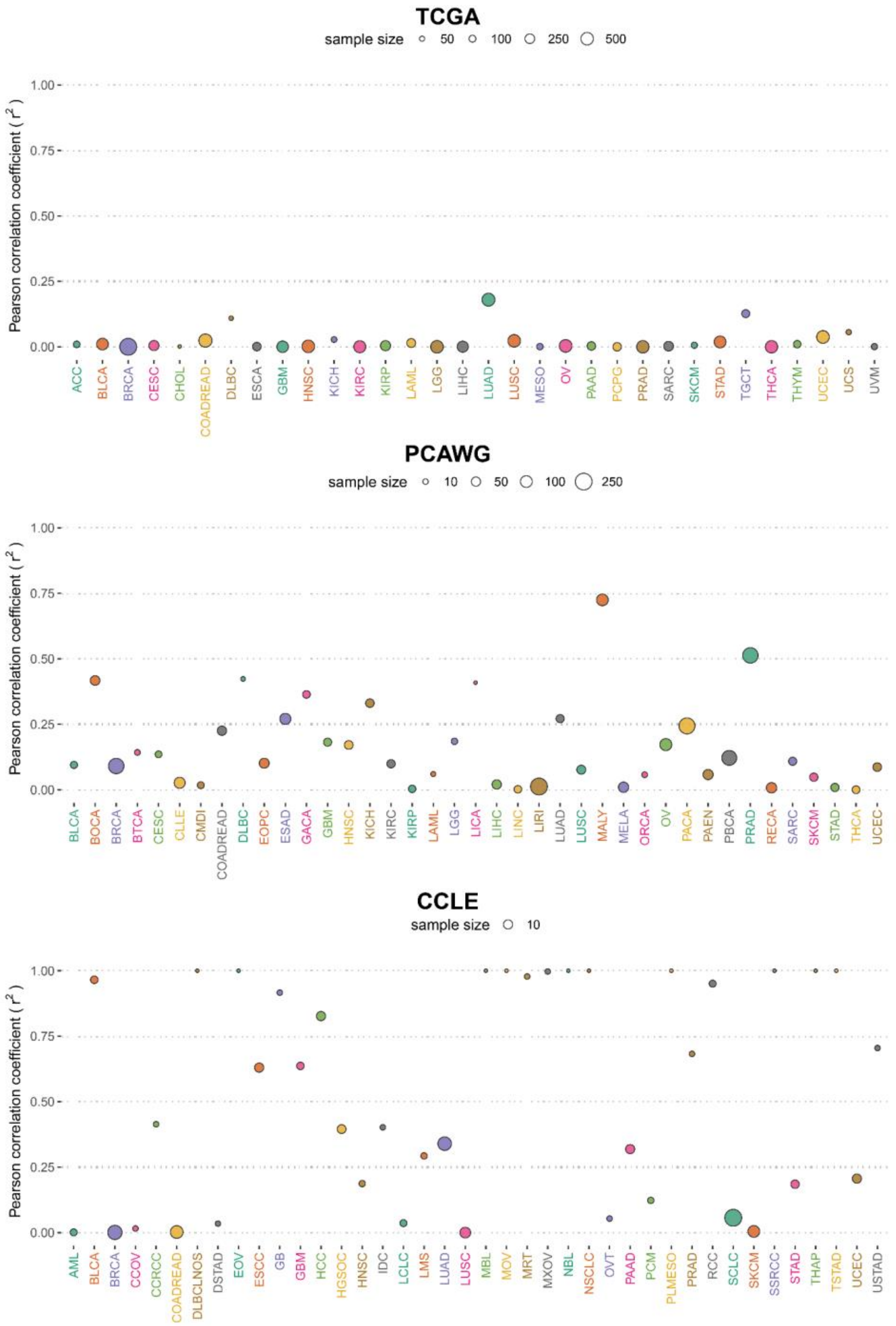
TCGA
sample size 50 100 250 500
Pearson correlation coefficient ( r² )
1.00
0.75
0.50
0.25
0.00
ACC BLCA BRCA CESC CHOL COADREAD DLBC ESCA GBM HNSC KICH KIRC KIRP LAML LGG LIHC LUAD LUSC MESO OV PAAD PCPG PRAD SARC SKCM STAD TGCT THCA THYM UCEC UCS UVM
PCAWG
sample size 10 50 100 250
Pearson correlation coefficient ( r² )
1.00
0.75
0.50
0.25
0.00
BLCA BOCA BRCA BTCA CESC CLLE CMDI COADREAD DLBC EOPC ESAD GACA GBM HNSC KICH KIRC KIRP LAML LGG LICA LIHC LINC LIRI LUAD LUSC MALY MELA ORCA OV PACA PAEN PBCA PRAD RECA SARC SKCM STAD THCA UCEC
CCLE
sample size 10
Pearson correlation coefficient ( r² )
1.00
0.75
0.50
0.25
0.00
AML BLCA BRCA CCOV CCRCC COADREAD DLBCLNOS DSTAD EOV ESCC GB GBM HCC HGSOC HNSC IDC LCLC LMS LUAD LUSC MBL MOV MRT MXOV NBL NSCLC OVT PAAD PCM PLMESO PRAD RCC SCLC SKCM SSRCC STAD THAP TSTAD UCEC USTAD

**Figure S6. TBL and TMB are poorly correlated in most cancer types.** Pan-cancer overview of the correlation between the TBL and the TMB reported with the Pearson correlation coefficient ($r^2$) for data from primary cancer (PCAWG and TCGA) and cell lines (CCLE). The size of the dot indicates the sample size. Color marks the type of cancer. Tumor type abbreviations from the CCLE cell lines were retrieved from OncoTree. ACC: Acute Myeloid Leukemia, BLCA: Bladder Urothelial Carcinoma, BRCA: Breast invasive carcinoma, CESC: Cervical squamous cell carcinoma and endocervical adenocarcinoma, CHOL: Cholangiocarcinoma, COADREAD: Colon and Rectum adenocarcinoma, DLBC: Lymphoid Neoplasm Diffuse Large B-cell Lymphoma , ESCA: Esophageal carcinoma, GBM: Glioblastoma multiforme, HNSC: Head and Neck squamous cell carcinoma, KICH: Kidney Chromophobe , KIRC: Kidney renal clear cell carcinoma, KIRP: Kidney renal papillary cell carcinoma, LAML: Acute Myeloid Leukemia, LGG: Brain Lower Grade Glioma, LIHC: Liver hepatocellular carcinoma, LUAD: Lung adenocarcinoma, LUSC: Lung squamous cell carcinoma, MESO: Mesothelioma , OV: Ovarian serous cystadenocarcinoma, PAAD: Pancreatic adenocarcinoma, PCPG: Pheochromocytoma and Paraganglioma, PRAD: Prostate adenocarcinoma, SARC: Sarcoma, SKCM: Skin Cutaneous Melanoma, STAD: Stomach adenocarcinoma, TGCT: Testicular Germ Cell Tumors, THCA: Thyroid carcinoma, THYM: Thymoma, UCEC: Uterine Corpus Endometrial Carcinoma, UCS: Uterine Carcinosarcoma, UVM: Uveal Melanoma. BLCA: Bladder Urothelial Carcinoma, BOCA: Bone Cancer, BTCA: Biliary tract cancer, CLLE: Chronic Lymphocytic Leukemia, CMDI: Chronic Myeloid Disorders, DLBC: Lymphoid Neoplasm Diffuse Large B-cell Lymphoma, EOPC: Early Onset Prostate Cancer, ESAD: Esophageal Adenocarcinoma, GACA: Gastric Cancer, KICH: Kidney Chromophobe, LICA: Liver Cancer, LINC: Liver Cancer, LIRI: Liver Cancer, MALY: Malignant Lymphoma, MELA: Melanoma, ORCA: Oral Cancer OV: Ovarian serous cystadenocarcinoma, PACA: Pancreatic Cancer, PAEN: Pancreatic Cancer Endocrine neoplasms, PBCA: Pediatric Brain Cancer, RECA: Renal Cancer, UCEC: Uterine Corpus Endometrial Carcinoma.

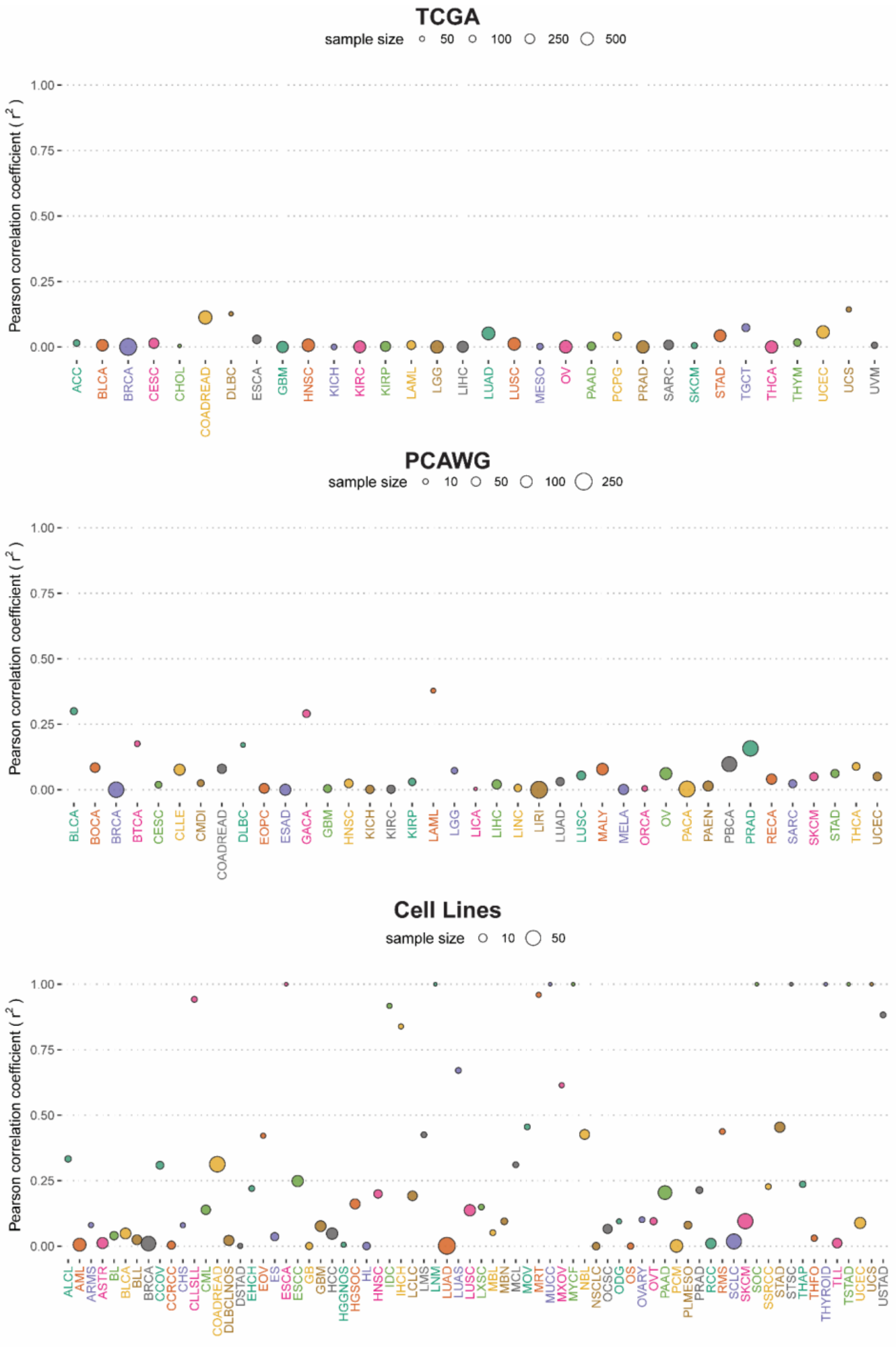
TCGA
sample size 50 100 250 500
Pearson correlation coefficient ( r² )
1.00
0.75
0.50
0.25
0.00
ACC BLCA BRCA CESC CHOL COADREAD DLBC ESCA GBM HNSC KICH KIRC KIRP LAML LGG LIHC LUAD LUSC MESO OV PAAD PCPG PRAD SARC SKCM STAD TGCT THCA THYM UCEC UCS UVM
PCAWG
sample size 10 50 100 250
Pearson correlation coefficient ( r² )
1.00
0.75
0.50
0.25
0.00
BLCA BOCA BRCA BTCA CESC CLLE CMDI COADREAD DLBC EOPC ESAD GACA GBM HNSC KICH KIRC KIRP LAML LGG LICA LIHC LINC LIRI LUAD LUSC MALY MELA ORCA OV PACA PAEN PBCA PRAD RECA SARC SKCM STAD THCA UCEC
Cell Lines
sample size 10 50
Pearson correlation coefficient ( r² )
1.00
0.75
0.50
0.25
0.00
ALCL AML ARMS ASTR BL BLCA BLL BRCA CCOV CCRCC CHS CLLSLL CML COADREAD DLBCLNOS DSTAD EHCH EOV ES ESCA ESCC GB GBM HCC HGGNOS HGSOC HL HNSC IDC IHCH LCLC LMS LNM LUAD LUAS LUSC LXSC MBL MBN MCL MOV MRT MUCC MXOV MYCF NBL NSCLC OCSC ODG OS OVARY OVT PAAD PCM PLMESO PRAD RCC RMS SCLC SKCM SOC SSRC STAD STSC THAP THFO THYROID TLL TSTAD UCEC UCS USTAD

**Figure S7. FGA and TMB are poorly correlated in most cancer types.** Pan-cancer overview of the correlation between FGA and TMB reported with Pearson's correlation coefficient ($r^2$) for data from primary cancer (PCAWG and TCGA) and cell lines (CCLE). The size of the dot indicates the sample size. Color marks the type of cancer. Tumor type abbreviations from the CCLE cell lines were retrieved from OncoTree. ACC: Acute Myeloid Leukemia, BLCA: Bladder Urothelial Carcinoma, BRCA: Breast invasive carcinoma, CESC: Cervical squamous cell carcinoma and endocervical adenocarcinoma, CHOL: Cholangiocarcinoma, COADREAD: Colon and Rectum adenocarcinoma, DLBC: Lymphoid Neoplasm Diffuse Large B-cell Lymphoma , ESCA: Esophageal carcinoma, GBM: Glioblastoma multiforme, HNSC: Head and Neck squamous cell carcinoma, KICH: Kidney Chromophobe , KIRC: Kidney renal clear cell carcinoma, KIRP: Kidney renal papillary cell carcinoma, LAML: Acute Myeloid Leukemia, LGG: Brain Lower Grade Glioma, LIHC: Liver hepatocellular carcinoma, LUAD: Lung adenocarcinoma, LUSC: Lung squamous cell carcinoma, MESO: Mesothelioma , OV: Ovarian serous cystadenocarcinoma, PAAD: Pancreatic adenocarcinoma, PCPG: Pheochromocytoma and Paraganglioma, PRAD: Prostate adenocarcinoma, SARC: Sarcoma, SKCM: Skin Cutaneous Melanoma, STAD: Stomach adenocarcinoma, TGCT: Testicular Germ Cell Tumors, THCA: Thyroid carcinoma, THYM: Thymoma, UCEC: Uterine Corpus Endometrial Carcinoma, UCS: Uterine Carcinosarcoma, UVM: Uveal Melanoma. BLCA: Bladder Urothelial Carcinoma, BOCA: Bone Cancer, BTCA: Biliary tract cancer, CLLE: Chronic Lymphocytic Leukemia, CMDI: Chronic Myeloid Disorders, DLBC: Lymphoid Neoplasm Diffuse Large B-cell Lymphoma, EOPC: Early Onset Prostate Cancer, ESAD: Esophageal Adenocarcinoma, GACA: Gastric Cancer, KICH: Kidney Chromophobe, LICA: Liver Cancer, LINC: Liver Cancer, LIRI: Liver Cancer, MALY: Malignant Lymphoma, MELA: Melanoma, ORCA: Oral Cancer  OV: Ovarian serous cystadenocarcinoma, PACA: Pancreatic Cancer, PAEN: Pancreatic Cancer Endocrine neoplasms, PBCA: Pediatric Brain Cancer, RECA: Renal Cancer, UCEC: Uterine Corpus Endometrial Carcinoma.

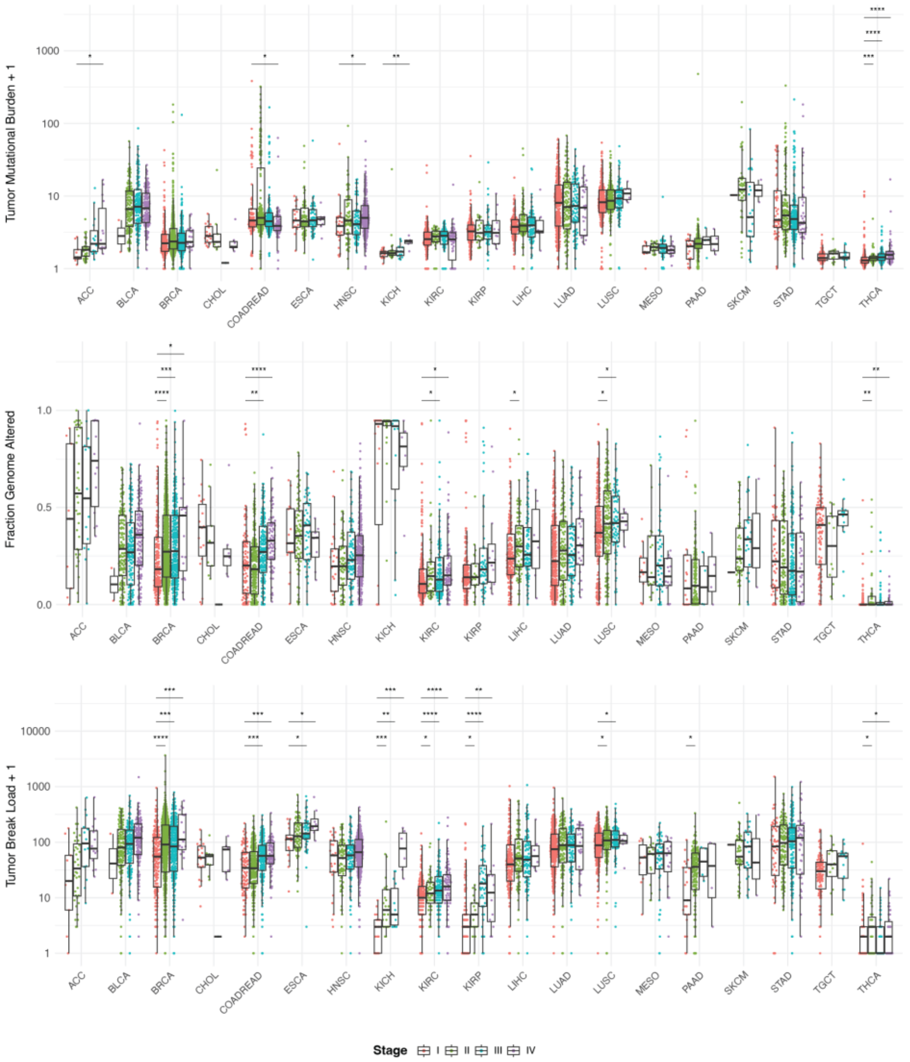


**Figure S8 Stage is associated with an increase in genomic instability measures.** Pan-cancer overview of the distribution of TMB (top), FGA (middle), and TBL (bottom) in TCGA across the different tumor stages (I-IV). TBL shows an increase in later stages in most cancer types. A Mann-Whitney U test was used to assess significance. Multiple hypothesis testing was performed using the Benjamini-Hochberg method. Significance levels: **** = $p_{adj}$ <0.0001, *** = $p_{adj}$ <0.001, ** = $p_{adj}$ <0.01, * = $p_{adj}$ <0.05. Comparisons without significance labels are not significant. ACC: Acute Myeloid Leukemia, BLCA: Bladder Urothelial Carcinoma, BRCA: Breast invasive carcinoma, CESC: Cervical squamous cell carcinoma and endocervical adenocarcinoma, CHOL: Cholangiocarcinoma, COADREAD: Colon and Rectum adenocarcinoma, DLBC: Lymphoid Neoplasm Diffuse Large B-cell Lymphoma , ESCA: Esophageal carcinoma, GBM: Glioblastoma multiforme, HNSC: Head and Neck squamous cell carcinoma, KICH: Kidney Chromophobe , KIRC: Kidney renal clear cell carcinoma, KIRP: Kidney renal papillary cell carcinoma, LAML: Acute Myeloid Leukemia, LGG: Brain Lower Grade Glioma, LIHC: Liver hepatocellular carcinoma, LUAD: Lung adenocarcinoma, LUSC: Lung squamous cell carcinoma, MESO: Mesothelioma , OV: Ovarian serous cystadenocarcinoma, PAAD: Pancreatic adenocarcinoma, PCPG: Pheochromocytoma and Paraganglioma, PRAD: Prostate adenocarcinoma, SARC: Sarcoma, SKCM: Skin Cutaneous Melanoma, STAD: Stomach adenocarcinoma, TGCT: Testicular Germ Cell Tumors, THCA: Thyroid carcinoma, THYM: Thymoma, UCEC: Uterine Corpus Endometrial Carcinoma, UCS: Uterine

Carcinosarcoma, UVM: Uveal Melanoma. BLCA: Bladder Urothelial Carcinoma, BOCA: Bone Cancer, BTCA: Biliary tract cancer, CLLE: Chronic Lymphocytic Leukemia, CMDI: Chronic Myeloid Disorders, DLBC: Lymphoid Neoplasm Diffuse Large B-cell Lymphoma, EOPC: Early Onset Prostate Cancer, ESAD: Esophageal Adenocarcinoma, GACA: Gastric Cancer, KICH: Kidney Chromophobe, LICA: Liver Cancer, LINC: Liver Cancer, LIRI: Liver Cancer, MALY: Malignant Lymphoma, MELA: Melanoma, ORCA: Oral Cancer  OV: Ovarian serous cystadenocarcinoma, PACA: Pancreatic Cancer, PAEN: Pancreatic Cancer Endocrine neoplasms, PBCA: Pediatric Brain Cancer, RECA: Renal Cancer, UCEC: Uterine Corpus Endometrial Carcinoma.

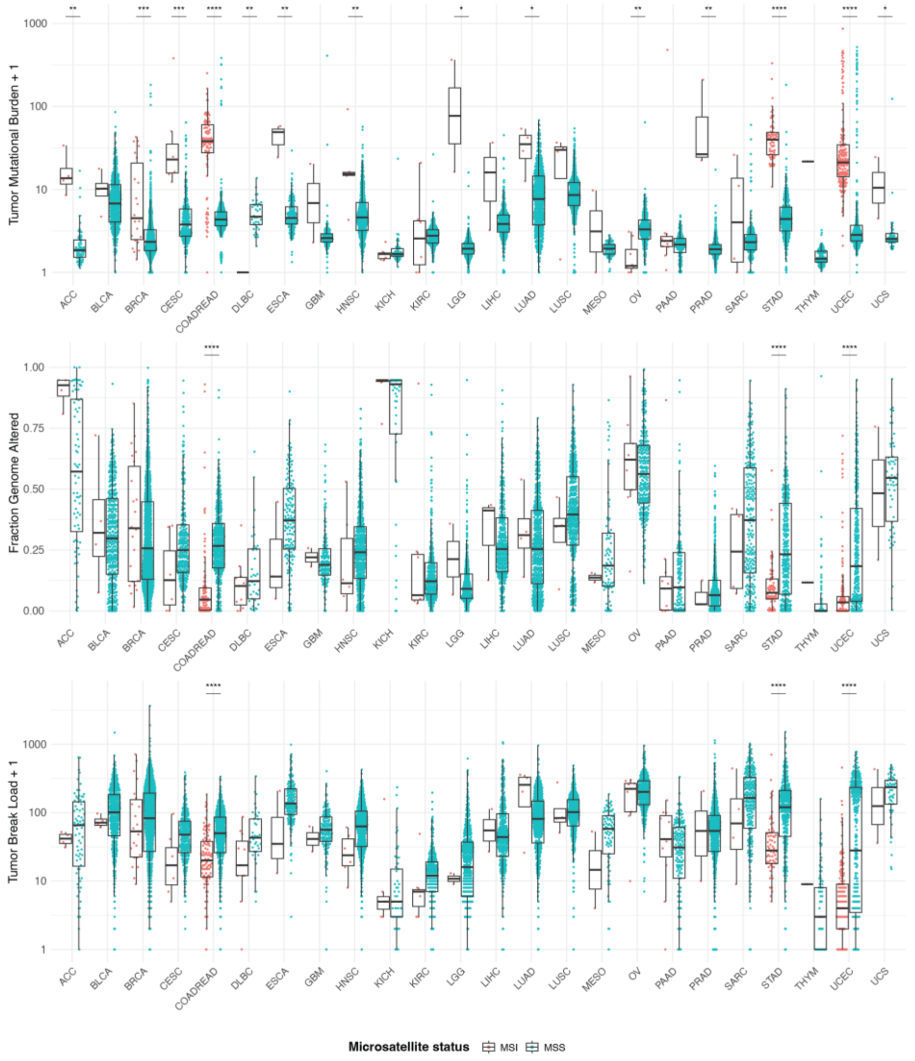


**Figure S9. The MSI phenotype is associated with differences in genomic instability measures measures.** Pan-cancer overview of the distribution of TMB (top), FGA (middle), and TBL (bottom) for MSI (red) and MSS (blue) tumor samples using data from the TCGA. MSI samples display a higher TMB in most cancer types. Some cancer types show a significantly higher TBL for MSS samples. A Mann-Whitney U test was used to assess significance. Multiple hypothesis testing was performed using the Benjamini-Hochberg method. Significance levels: **** = $p_{adj}$ <0.0001, *** = $p_{adj}$ <0.001, ** = $p_{adj}$ <0.01, * = $p_{adj}$ <0.05. Comparisons without significance labels are not significant.  ACC: Acute Myeloid Leukemia, BLCA: Bladder Urothelial Carcinoma, BRCA: Breast invasive carcinoma, CESC: Cervical squamous cell carcinoma and endocervical adenocarcinoma, CHOL: Cholangiocarcinoma, COADREAD: Colon and Rectum adenocarcinoma, DLBC: Lymphoid Neoplasm Diffuse Large B-cell Lymphoma , ESCA: Esophageal carcinoma, GBM: Glioblastoma multiforme, HNSC: Head and Neck squamous cell carcinoma, KICH: Kidney Chromophobe , KIRC: Kidney renal clear cell carcinoma, KIRP: Kidney renal papillary cell carcinoma, LAML: Acute Myeloid Leukemia, LGG: Brain Lower Grade Glioma, LIHC: Liver hepatocellular carcinoma, LUAD: Lung adenocarcinoma, LUSC: Lung squamous cell carcinoma, MESO: Mesothelioma , OV: Ovarian serous cystadenocarcinoma, PAAD: Pancreatic adenocarcinoma, PCPG: Pheochromocytoma and Paraganglioma, PRAD: Prostate adenocarcinoma, SARC: Sarcoma, SKCM: Skin Cutaneous Melanoma, STAD: Stomach adenocarcinoma, TGCT: Testicular Germ Cell Tumors, THCA: Thyroid carcinoma, THYM: Thymoma, UCEC: Uterine Corpus Endometrial Carcinoma, UCS: Uterine Carcinosarcoma, UVM: Uveal Melanoma. BLCA: Bladder Urothelial Carcinoma, BOCA: Bone Cancer, BTCA: Biliary tract cancer, CLLE: Chronic Lymphocytic Leukemia, CMDI: Chronic Myeloid Disorders, DLBC: Lymphoid Neoplasm Diffuse Large B-cell Lymphoma, EOPC: Early Onset Prostate Cancer, ESAD: Esophageal Adenocarcinoma, GACA: Gastric Cancer, KICH: Kidney Chromophobe, LICA: Liver Cancer, LINC: Liver Cancer, LIRI: Liver Cancer, MALY: Malignant Lymphoma, MELA: Melanoma, ORCA: Oral Cancer  OV: Ovarian serous cystadenocarcinoma, PACA: Pancreatic Cancer, PAEN: Pancreatic Cancer Endocrine neoplasms, PBCA: Pediatric Brain Cancer, RECA: Renal Cancer, UCEC: Uterine Corpus Endometrial Carcinoma.

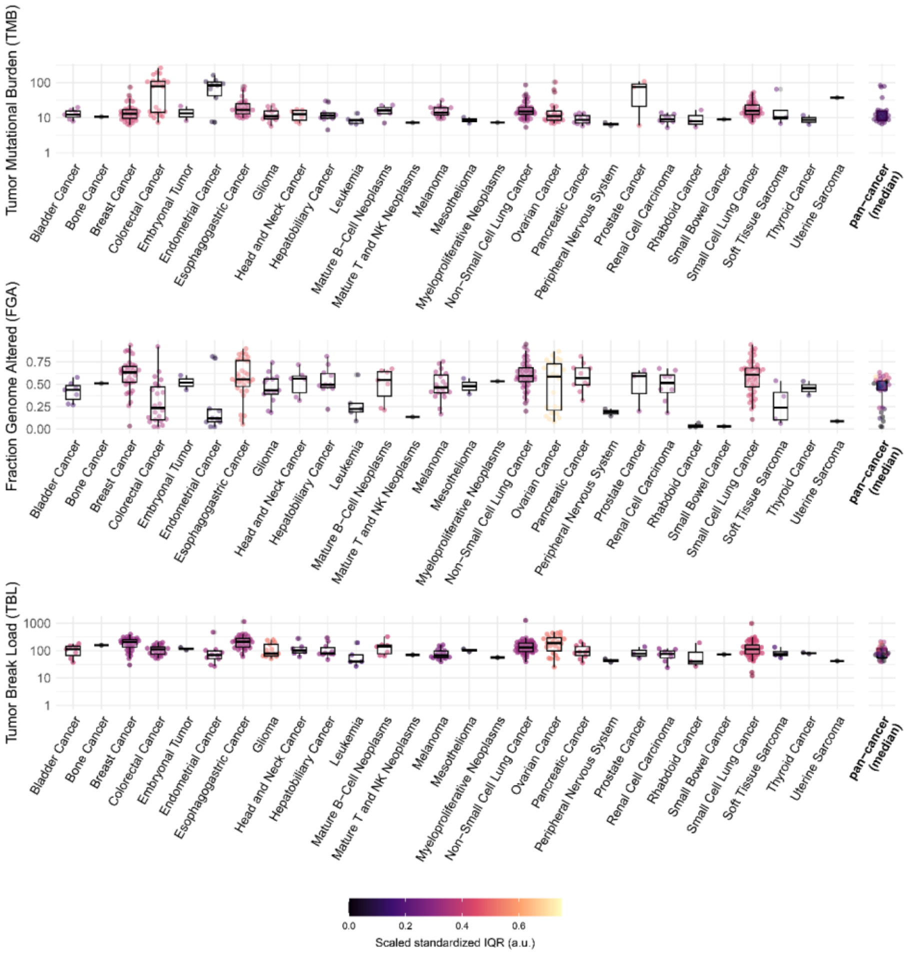


**Figure S10**. **High levels of genomic instability in cancer cell lines.** Pan-cancer overview of the tumor mutational burden (TMB), the fraction genome altered (FGA), and the tumor break load (TBL) in cancer cell lines. TMB and TBL are visualized on a $\log_{10}$ scale. The color indicates the scaled standardized interquartile range (IQR) calculated from the ranked measures. The medians of all cancer types are reported as pan-cancer (median).

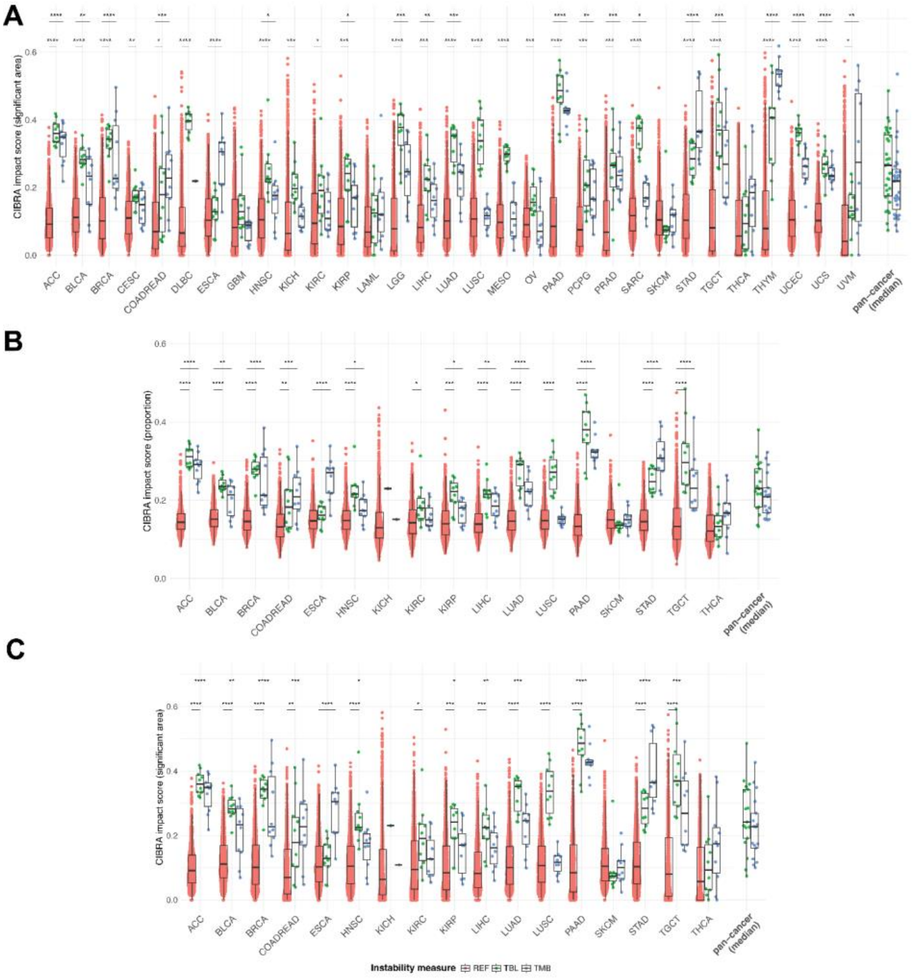


**Figure S11 TBL is associated with altered gene expression in most cancer types using data from TCGA.** CIBRA impact score (significant area) for comparison of tumor samples from both extremes of the genomic instability distribution. For each genomic instability measure and cancer type, we dichotomized the samples into a high and low group (top and bottom 25%), creating 10 subsample datasets of 10 HIGH and 10 LOW samples each. The resulting CIBRA scores were compared to a reference distribution of scores from 1000 random permutations of the dataset. We performed the analysis for all primary cancer samples (**A**), as well as exclusively MSS samples in stage I-II (**B, C**). Cancer types for which no stage or MSS/MSI information was available were filtered out for the second analysis. We report two impact scores: CIBRA proportion (**B**) and significant area (**A, C**). Statistical significance was assessed using a Mann-Whitney U test. We used multiple hypothesis correction using the Benjamini-Hochberg method. Significance levels: **** = $p_{adj} < 0.0001$, *** = $p_{adj} < 0.001$, ** = $p_{adj} < 0.01$, * = $p_{adj} < 0.05$. Comparisons without significance labels are not significant. ACC: Acute Myeloid Leukemia, BLCA: Bladder Urothelial Carcinoma, BRCA: Breast invasive carcinoma, CESC: Cervical squamous cell carcinoma and endocervical adenocarcinoma, CHOL: Cholangiocarcinoma, COADREAD: Colon and Rectum adenocarcinoma, DLBC: Lymphoid Neoplasm Diffuse Large B-cell Lymphoma , ESCA: Esophageal carcinoma, GBM: Glioblastoma multiforme, HNSC: Head and Neck squamous cell carcinoma, KICH: Kidney Chromophobe , KIRC: Kidney renal clear cell carcinoma, KIRP: Kidney renal papillary cell carcinoma, LAML: Acute Myeloid Leukemia, LGG: Brain Lower Grade Glioma, LIHC: Liver hepatocellular carcinoma, LUAD: Lung adenocarcinoma, LUSC: Lung squamous cell carcinoma, MESO: Mesothelioma , OV: Ovarian serous cystadenocarcinoma, PAAD: Pancreatic adenocarcinoma, PCPG: Pheochromocytoma and Paraganglioma, PRAD: Prostate adenocarcinoma, SARC: Sarcoma, SKCM: Skin Cutaneous Melanoma, STAD: Stomach adenocarcinoma, TGCT: Testicular Germ Cell Tumors, THCA: Thyroid carcinoma, THYM: Thymoma, UCEC: Uterine Corpus Endometrial Carcinoma, UCS: Uterine

Carcinosarcoma, UVM: Uveal Melanoma. BLCA: Bladder Urothelial Carcinoma, BOCA: Bone Cancer, BTCA: Biliary tract cancer, CLLE: Chronic Lymphocytic Leukemia, CMDI: Chronic Myeloid Disorders, DLBC: Lymphoid Neoplasm Diffuse Large B-cell Lymphoma, EOPC: Early Onset Prostate Cancer, ESAD: Esophageal Adenocarcinoma, GACA: Gastric Cancer, KICH: Kidney Chromophobe, LICA: Liver Cancer, LINC: Liver Cancer, LIRI: Liver Cancer, MALY: Malignant Lymphoma, MELA: Melanoma, ORCA: Oral Cancer  OV: Ovarian serous cystadenocarcinoma, PACA: Pancreatic Cancer, PAEN: Pancreatic Cancer Endocrine neoplasms, PBCA: Pediatric Brain Cancer, RECA: Renal Cancer, UCEC: Uterine Corpus Endometrial Carcinoma.

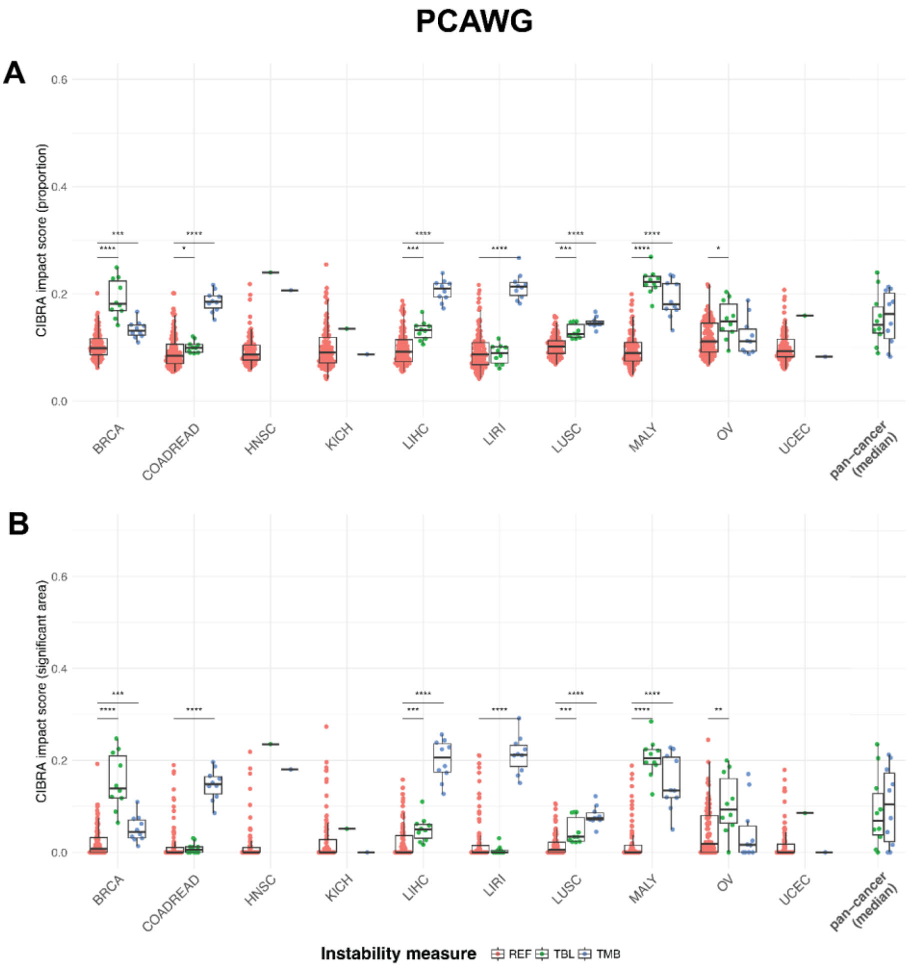


**Figure S12 CIBRA impact scores for the PCAWG dataset.** CIBRA impact score (significant area) for the comparison of tumor samples from both extremes of the genomic instability distribution. For each genomic instability measure and cancer type, we dichotomized the samples into a high and low group (top and bottom 25%), creating 10 subsample datasets of 10 HIGH and 10 LOW samples each. The resulting CIBRA scores were compared to a reference distribution of 100 random permutations of the dataset. We report two impact scores: CIBRA proportion (**A**) and significant area (**B**). Statistical significance was assessed using a Mann-Whitney U test. We used multiple hypothesis correction using the Benjamini-Hochberg method. Significance levels: **** = $p_{adj}$ <0.0001, *** = $p_{adj}$ <0.001, ** = $p_{adj}$ <0.01, * = $p_{adj}$ <0.05. Comparisons without significance labels are not significant. ACC: Acute Myeloid Leukemia, BLCA: Bladder Urothelial Carcinoma, BRCA: Breast invasive carcinoma, CESC: Cervical squamous cell carcinoma and endocervical adenocarcinoma, CHOL: Cholangiocarcinoma, COADREAD: Colon and Rectum adenocarcinoma, DLBC: Lymphoid Neoplasm Diffuse Large B-cell Lymphoma , ESCA: Esophageal carcinoma, GBM: Glioblastoma multiforme, HNSC: Head and Neck squamous cell carcinoma, KICH: Kidney Chromophobe , KIRC: Kidney renal clear cell carcinoma, KIRP: Kidney renal papillary cell carcinoma, LAML: Acute Myeloid Leukemia, LGG: Brain Lower Grade Glioma, LIHC: Liver hepatocellular carcinoma, LUAD: Lung adenocarcinoma, LUSC: Lung squamous cell carcinoma, MESO: Mesothelioma , OV: Ovarian serous cystadenocarcinoma, PAAD: Pancreatic adenocarcinoma, PCPG: Pheochromocytoma and Paraganglioma, PRAD: Prostate adenocarcinoma, SARC: Sarcoma, SKCM: Skin Cutaneous Melanoma, STAD: Stomach adenocarcinoma, TGCT: Testicular Germ Cell Tumors, THCA: Thyroid carcinoma, THYM: Thymoma, UCEC: Uterine Corpus Endometrial Carcinoma, UCS: Uterine Carcinosarcoma, UVM: Uveal Melanoma. BLCA: Bladder Urothelial Carcinoma, BOCA: Bone Cancer, BTCA: Biliary tract cancer, CLLE: Chronic Lymphocytic Leukemia, CMDI: Chronic Myeloid Disorders, DLBC: Lymphoid Neoplasm Diffuse Large B-cell Lymphoma, EOPC:

Early Onset Prostate Cancer, ESAD: Esophageal Adenocarcinoma, GACA: Gastric Cancer, KICH: Kidney Chromophobe, LICA: Liver Cancer, LINC: Liver Cancer, LIRI: Liver Cancer, MALY: Malignant Lymphoma, MELA: Melanoma, ORCA: Oral Cancer OV: Ovarian serous cystadenocarcinoma, PACA: Pancreatic Cancer, PAEN: Pancreatic Cancer Endocrine neoplasms, PBCA: Pediatric Brain Cancer, RECA: Renal Cancer, UCEC: Uterine Corpus Endometrial Carcinoma.

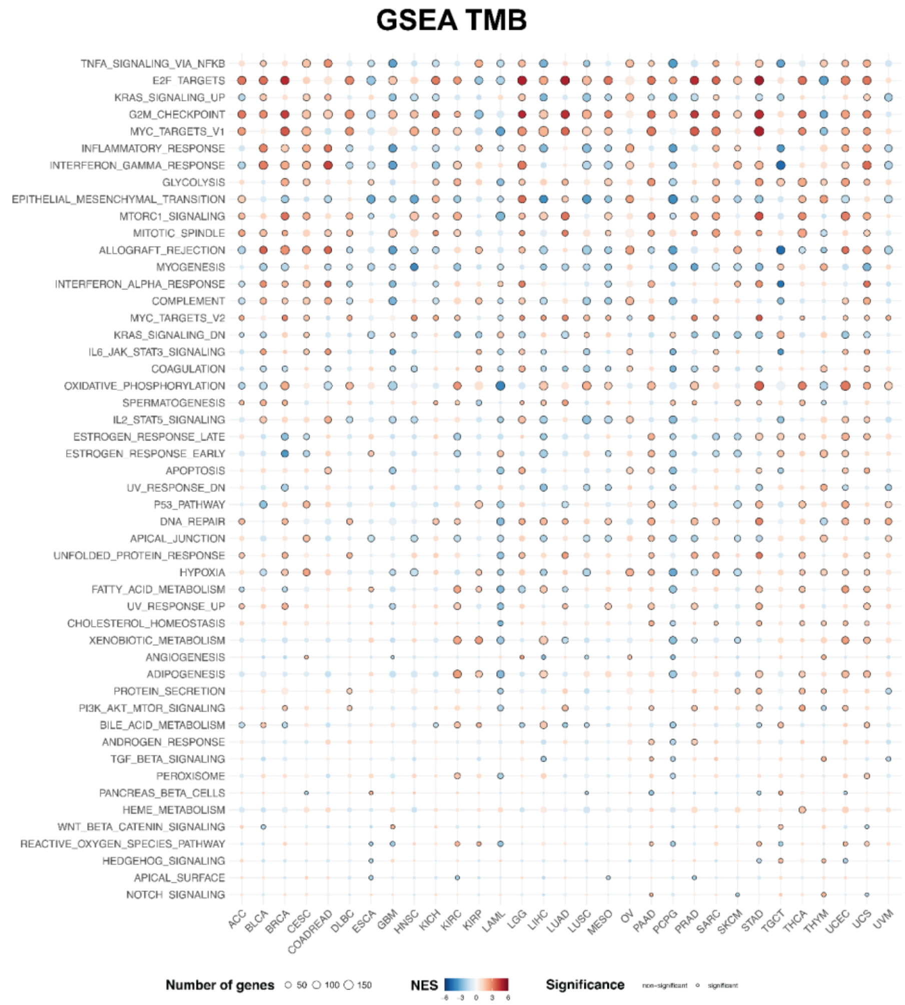


**Figure S13. Results of GSEA for TMB-high vs TMB-low using the MSigDb Hallmark Gene Set.** Pathways are ranked according to the number of cancer types in which they are differentially expressed in the TBL-high vs TBL-low comparison. Colors indicate Normalized Enrichment Scores (NSE; red for up-regulated pathways, blue for down-regulated pathways). Multiple testing correction was performed using the Benjamini-Hochberg method. Black circles indicate significance (adjusted p-value < 0.01). ACC: Acute Myeloid Leukemia, BLCA: Bladder Urothelial Carcinoma, BRCA: Breast invasive carcinoma, CESC: Cervical squamous cell carcinoma and endocervical adenocarcinoma, CHOL: Cholangiocarcinoma, COADREAD: Colon and Rectum adenocarcinoma, DLBC: Lymphoid Neoplasm Diffuse Large B-cell Lymphoma , ESCA: Esophageal carcinoma, GBM: Glioblastoma multiforme, HNSC: Head and Neck squamous cell carcinoma, KICH: Kidney Chromophobe , KIRC: Kidney renal clear cell carcinoma, KIRP: Kidney renal papillary cell carcinoma, LAML: Acute Myeloid Leukemia, LGG: Brain Lower Grade Glioma, LIHC: Liver hepatocellular carcinoma, LUAD: Lung adenocarcinoma, LUSC: Lung squamous cell carcinoma, MESO: Mesothelioma , OV: Ovarian serous cystadenocarcinoma, PAAD: Pancreatic adenocarcinoma, PCPG: Pheochromocytoma and Paraganglioma, PRAD: Prostate adenocarcinoma, SARC: Sarcoma, SKCM: Skin Cutaneous Melanoma, STAD: Stomach adenocarcinoma, TGCT: Testicular Germ Cell Tumors, THCA: Thyroid carcinoma, THYM: Thymoma, UCEC: Uterine Corpus Endometrial Carcinoma, UCS: Uterine Carcinosarcoma, UVM: Uveal Melanoma. BLCA: Bladder Urothelial Carcinoma, BOCA: Bone Cancer, BTCA: Biliary tract cancer, CLLE: Chronic Lymphocytic Leukemia, CMDI: Chronic Myeloid Disorders, DLBC: Lymphoid Neoplasm Diffuse Large B-cell Lymphoma, EOPC: Early Onset Prostate Cancer, ESAD: Esophageal Adenocarcinoma, GACA: Gastric Cancer, KICH: Kidney Chromophobe, LICA: Liver Cancer, LINC: Liver Cancer, LIRI: Liver Cancer, MALY: Malignant Lymphoma, MELA: Melanoma, ORCA: Oral Cancer  OV: Ovarian serous cystadenocarcinoma, PACA: Pancreatic Cancer, PAEN: Pancreatic Cancer Endocrine neoplasms, PBCA: Pediatric Brain Cancer, RECA: Renal Cancer, UCEC: Uterine Corpus Endometrial Carcinoma.

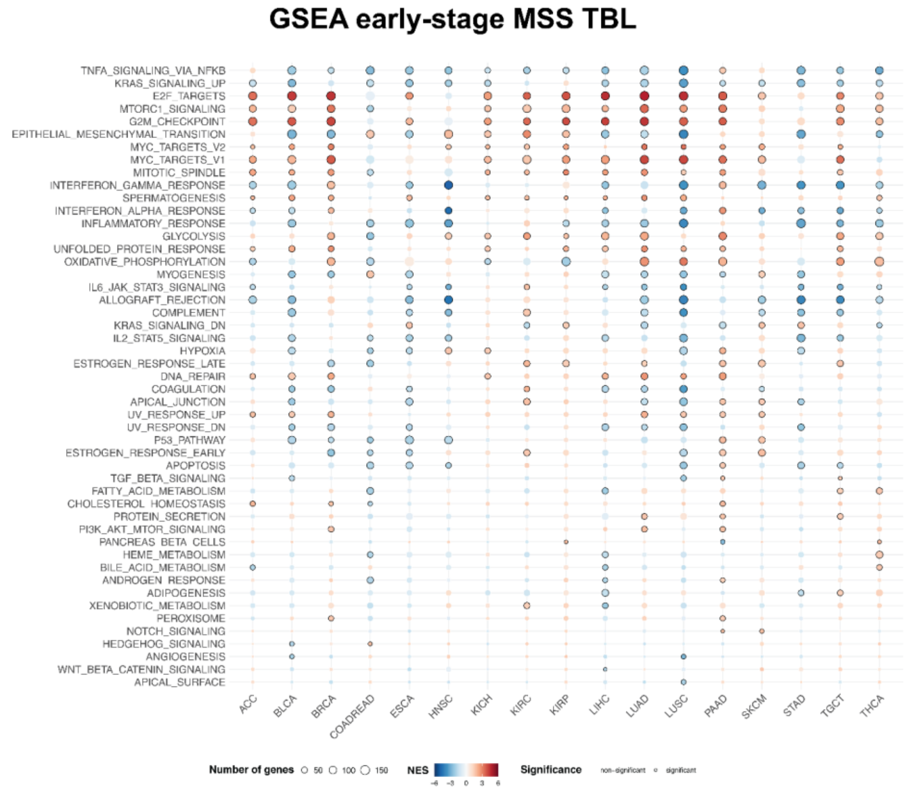


**Figure S14. Results of GSEA analysis for TBL-high vs TBL-low in early-stage MSS (TCGA) using the MSigDb Hallmark Gene Set.** Pathways are ordered based on the number of cancer types in which they are differentially expressed. Colors indicate Normalized Enrichment Scores (NES; red for up-regulated pathways, blue for down-regulated pathways). Multiple testing correction was performed using the Benjamini-Hochberg method. Black circles indicate significance (adjusted p-value < 0.01). ACC: Acute Myeloid Leukemia, BLCA: Bladder Urothelial Carcinoma, BRCA: Breast invasive carcinoma, CESC: Cervical squamous cell carcinoma and endocervical adenocarcinoma, CHOL: Cholangiocarcinoma, COADREAD: Colon and Rectum adenocarcinoma, DLBC: Lymphoid Neoplasm Diffuse Large B-cell Lymphoma , ESCA: Esophageal carcinoma, GBM: Glioblastoma multiforme, HNSC: Head and Neck squamous cell carcinoma, KICH: Kidney Chromophobe , KIRC: Kidney renal clear cell carcinoma, KIRP: Kidney renal papillary cell carcinoma, LAML: Acute Myeloid Leukemia, LGG: Brain Lower Grade Glioma, LIHC: Liver hepatocellular carcinoma, LUAD: Lung adenocarcinoma, LUSC: Lung squamous cell carcinoma, MESO: Mesothelioma , OV: Ovarian serous cystadenocarcinoma, PAAD: Pancreatic adenocarcinoma, PCPG: Pheochromocytoma and Paraganglioma, PRAD: Prostate adenocarcinoma, SARC: Sarcoma, SKCM: Skin Cutaneous Melanoma, STAD: Stomach adenocarcinoma, TGCT: Testicular Germ Cell Tumors, THCA: Thyroid carcinoma, THYM: Thymoma, UCEC: Uterine Corpus Endometrial Carcinoma, UCS: Uterine Carcinosarcoma, UVM: Uveal Melanoma. BLCA: Bladder Urothelial Carcinoma, BOCA: Bone Cancer, BTCA: Biliary tract cancer, CLLE: Chronic Lymphocytic Leukemia, CMDI: Chronic Myeloid Disorders, DLBC: Lymphoid Neoplasm Diffuse Large B-cell Lymphoma, EOPC: Early Onset Prostate Cancer, ESAD: Esophageal Adenocarcinoma, GACA: Gastric Cancer, KICH: Kidney Chromophobe, LICA: Liver Cancer, LINC: Liver Cancer, LIRI: Liver Cancer, MALY: Malignant Lymphoma, MELA: Melanoma, ORCA: Oral Cancer OV: Ovarian serous cystadenocarcinoma, PACA: Pancreatic Cancer, PAEN: Pancreatic Cancer Endocrine neoplasms, PBCA: Pediatric Brain Cancer, RECA: Renal Cancer, UCEC: Uterine Corpus Endometrial Carcinoma.

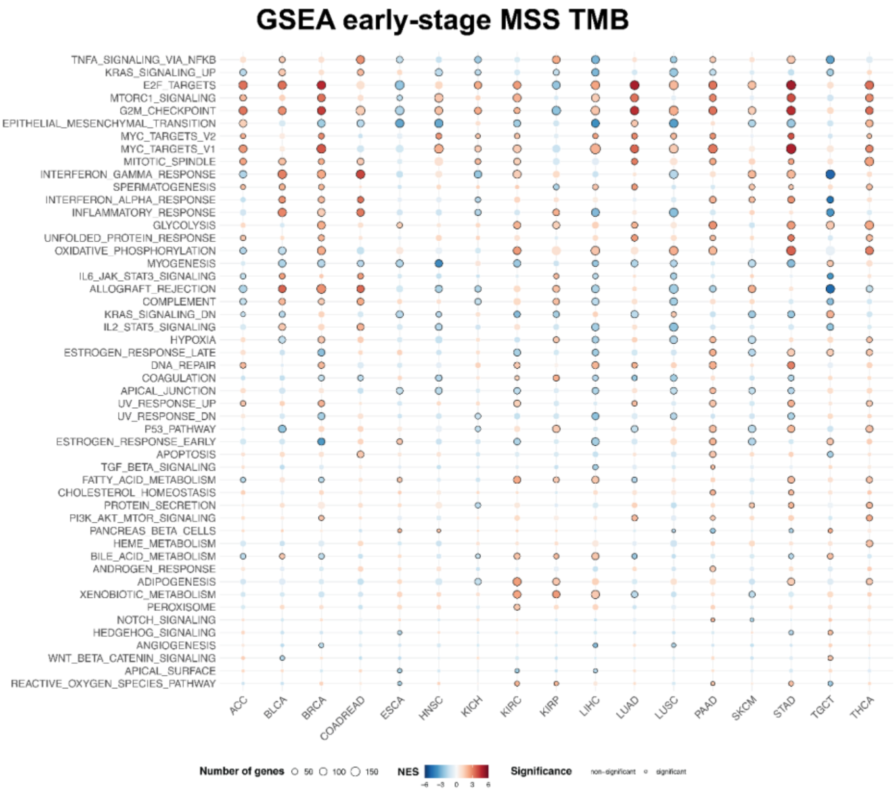


**Figure S15. Results of GSEA analysis for TMB-high vs TMB-low in early-stage MSS (TCGA) using the MSigDb Hallmark Gene Set.** Pathways are ordered based on the number of cancer types in which they are differentially expressed in the early-MSS TBL-high vs TBL-low comparison. Colors indicate Normalized Enrichment Scores (NES; red for up-regulated pathways, blue for down-regulated pathways). Multiple testing correction was performed using the Benjamini-Hochberg method. Black circles indicate significance (adjusted p-value < 0.01). ACC: Acute Myeloid Leukemia, BLCA: Bladder Urothelial Carcinoma, BRCA: Breast invasive carcinoma, CESC: Cervical squamous cell carcinoma and endocervical adenocarcinoma, CHOL: Cholangiocarcinoma, COADREAD: Colon and Rectum adenocarcinoma, DLBC: Lymphoid Neoplasm Diffuse Large B-cell Lymphoma , ESCA: Esophageal carcinoma, GBM: Glioblastoma multiforme, HNSC: Head and Neck squamous cell carcinoma, KICH: Kidney Chromophobe , KIRC: Kidney renal clear cell carcinoma, KIRP: Kidney renal papillary cell carcinoma, LAML: Acute Myeloid Leukemia, LGG: Brain Lower Grade Glioma, LIHC: Liver hepatocellular carcinoma, LUAD: Lung adenocarcinoma, LUSC: Lung squamous cell carcinoma, MESO: Mesothelioma , OV: Ovarian serous cystadenocarcinoma, PAAD: Pancreatic adenocarcinoma, PCPG: Pheochromocytoma and Paraganglioma, PRAD: Prostate adenocarcinoma, SARC: Sarcoma, SKCM: Skin Cutaneous Melanoma, STAD: Stomach adenocarcinoma, TGCT: Testicular Germ Cell Tumors, THCA: Thyroid carcinoma, THYM: Thymoma, UCEC: Uterine Corpus Endometrial Carcinoma, UCS: Uterine Carcinosarcoma, UVM: Uveal Melanoma. BLCA: Bladder Urothelial Carcinoma, BOCA: Bone Cancer, BTCA: Biliary tract cancer, CLLE: Chronic Lymphocytic Leukemia, CMDI: Chronic Myeloid Disorders, DLBC: Lymphoid Neoplasm Diffuse Large B-cell Lymphoma, EOPC: Early Onset Prostate Cancer, ESAD: Esophageal Adenocarcinoma, GACA: Gastric Cancer, KICH: Kidney Chromophobe, LICA: Liver Cancer, LINC: Liver Cancer, LIRI: Liver Cancer, MALY: Malignant Lymphoma, MELA: Melanoma, ORCA: Oral Cancer OV: Ovarian serous cystadenocarcinoma, PACA: Pancreatic Cancer, PAEN: Pancreatic Cancer Endocrine neoplasms, PBCA: Pediatric Brain Cancer, RECA: Renal Cancer, UCEC: Uterine Corpus Endometrial Carcinoma.

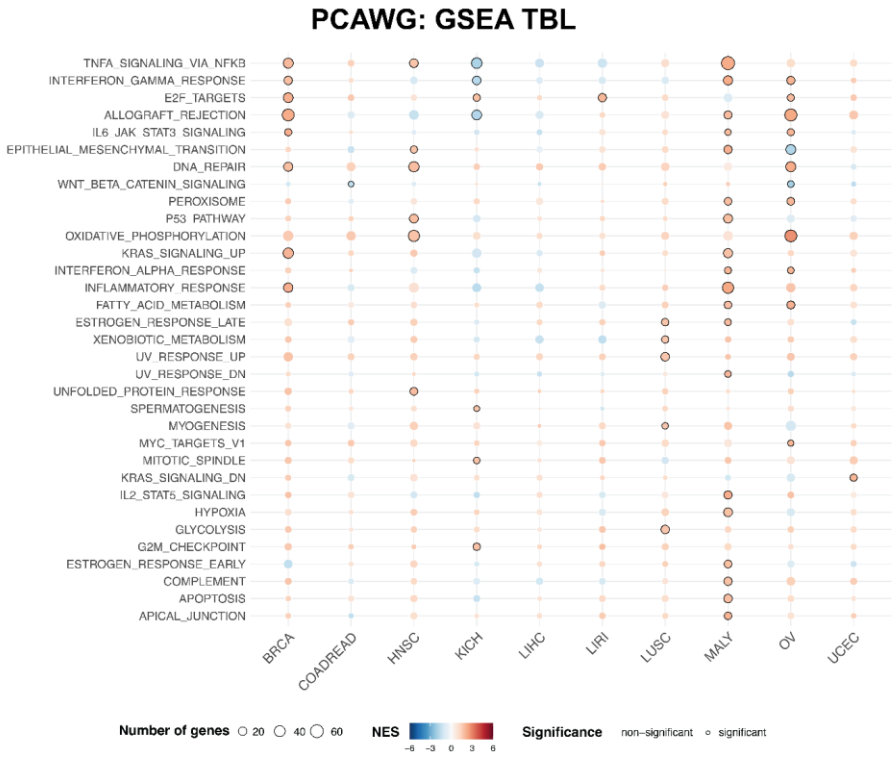


**Figure S16. Results of GSEA analysis for TBL-high vs TBL-low (PCAWG) using the MSigDb Hallmark Gene Set.** Pathways are ordered based on the number of cancer types in which they are differentially expressed. Colors indicate Normalized Enrichment Scores (NES; red for up-regulated pathways, blue for down-regulated pathways). Multiple testing correction was performed using the Benjamini-Hochberg method. Black circles indicate significance (adjusted p-value < 0.01). ACC: Acute Myeloid Leukemia, BLCA: Bladder Urothelial Carcinoma, BRCA: Breast invasive carcinoma, CESC: Cervical squamous cell carcinoma and endocervical adenocarcinoma, CHOL: Cholangiocarcinoma, COADREAD: Colon and Rectum adenocarcinoma, DLBC: Lymphoid Neoplasm Diffuse Large B-cell Lymphoma , ESCA: Esophageal carcinoma, GBM: Glioblastoma multiforme, HNSC: Head and Neck squamous cell carcinoma, KICH: Kidney Chromophobe , KIRC: Kidney renal clear cell carcinoma, KIRP: Kidney renal papillary cell carcinoma, LAML: Acute Myeloid Leukemia, LGG: Brain Lower Grade Glioma, LIHC: Liver hepatocellular carcinoma, LUAD: Lung adenocarcinoma, LUSC: Lung squamous cell carcinoma, MESO: Mesothelioma , OV: Ovarian serous cystadenocarcinoma, PAAD: Pancreatic adenocarcinoma, PCPG: Pheochromocytoma and Paraganglioma, PRAD: Prostate adenocarcinoma, SARC: Sarcoma, SKCM: Skin Cutaneous Melanoma, STAD: Stomach adenocarcinoma, TGCT: Testicular Germ Cell Tumors, THCA: Thyroid carcinoma, THYM: Thymoma, UCEC: Uterine Corpus Endometrial Carcinoma, UCS: Uterine Carcinosarcoma, UVM: Uveal Melanoma. BLCA: Bladder Urothelial Carcinoma, BOCA: Bone Cancer, BTCA: Biliary tract cancer, CLLE: Chronic Lymphocytic Leukemia, CMDI: Chronic Myeloid Disorders, DLBC: Lymphoid Neoplasm Diffuse Large B-cell Lymphoma, EOPC: Early Onset Prostate Cancer, ESAD: Esophageal Adenocarcinoma, GACA: Gastric Cancer, KICH: Kidney Chromophobe, LICA: Liver Cancer, LINC: Liver Cancer, LIRI: Liver Cancer, MALY: Malignant Lymphoma, MELA: Melanoma, ORCA: Oral Cancer OV: Ovarian serous cystadenocarcinoma, PACA: Pancreatic Cancer, PAEN: Pancreatic Cancer Endocrine neoplasms, PBCA: Pediatric Brain Cancer, RECA: Renal Cancer, UCEC: Uterine Corpus Endometrial Carcinoma.

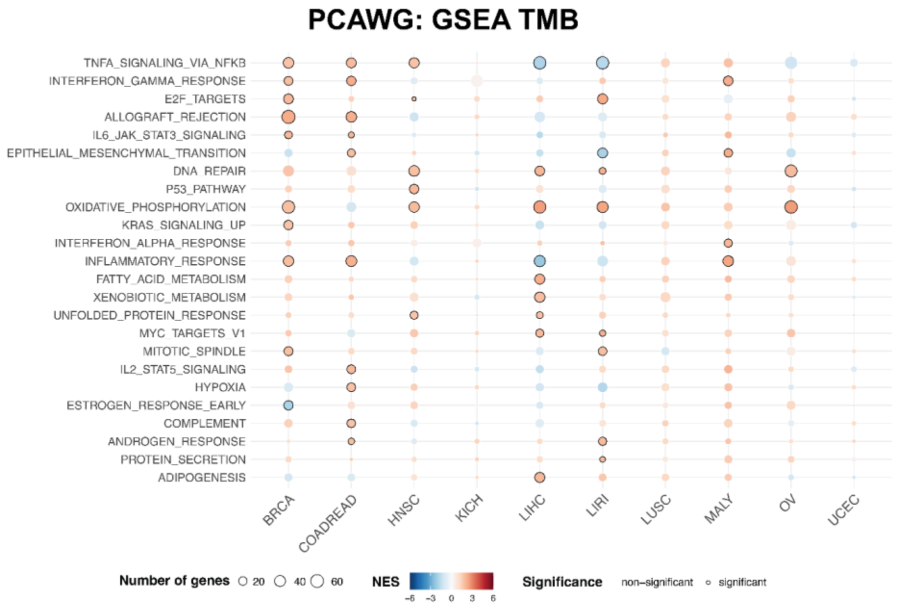


**Figure S17. Results of GSEA analysis for TMB-high vs TMB-low (PCAWG) using the MSigDb Hallmark Gene Set** Pathways are ordered based on the number of cancer types in which they are differentially expressed in the PCAWG TBL-high vs TBL-low comparison. Colors indicate Normalized Enrichment Scores (NES; red for up-regulated pathways, blue for down-regulated pathways). Multiple testing correction was performed using the Benjamini-Hochberg method. Black circles indicate significance (adjusted p-value < 0.01). ACC: Acute Myeloid Leukemia, BLCA: Bladder Urothelial Carcinoma, BRCA: Breast invasive carcinoma, CESC: Cervical squamous cell carcinoma and endocervical adenocarcinoma, CHOL: Cholangiocarcinoma, COADREAD: Colon and Rectum adenocarcinoma, DLBC: Lymphoid Neoplasm Diffuse Large B-cell Lymphoma , ESCA: Esophageal carcinoma, GBM: Glioblastoma multiforme, HNSC: Head and Neck squamous cell carcinoma, KICH: Kidney Chromophobe , KIRC: Kidney renal clear cell carcinoma, KIRP: Kidney renal papillary cell carcinoma, LAML: Acute Myeloid Leukemia, LGG: Brain Lower Grade Glioma, LIHC: Liver hepatocellular carcinoma, LUAD: Lung adenocarcinoma, LUSC: Lung squamous cell carcinoma, MESO: Mesothelioma , OV: Ovarian serous cystadenocarcinoma, PAAD: Pancreatic adenocarcinoma, PCPG: Pheochromocytoma and Paraganglioma, PRAD: Prostate adenocarcinoma, SARC: Sarcoma, SKCM: Skin Cutaneous Melanoma, STAD: Stomach adenocarcinoma, TGCT: Testicular Germ Cell Tumors, THCA: Thyroid carcinoma, THYM: Thymoma, UCEC: Uterine Corpus Endometrial Carcinoma, UCS: Uterine Carcinosarcoma, UVM: Uveal Melanoma. BLCA: Bladder Urothelial Carcinoma, BOCA: Bone Cancer, BTCA: Biliary tract cancer, CLLE: Chronic Lymphocytic Leukemia, CMDI: Chronic Myeloid Disorders, DLBC: Lymphoid Neoplasm Diffuse Large B-cell Lymphoma, EOPC: Early Onset Prostate Cancer, ESAD: Esophageal Adenocarcinoma, GACA: Gastric Cancer, KICH: Kidney Chromophobe, LICA: Liver Cancer, LINC: Liver Cancer, LIRI: Liver Cancer, MALY: Malignant Lymphoma, MELA: Melanoma, ORCA: Oral Cancer OV: Ovarian serous cystadenocarcinoma, PACA: Pancreatic Cancer, PAEN: Pancreatic Cancer Endocrine neoplasms, PBCA: Pediatric Brain Cancer, RECA: Renal Cancer, UCEC: Uterine Corpus Endometrial Carcinoma.

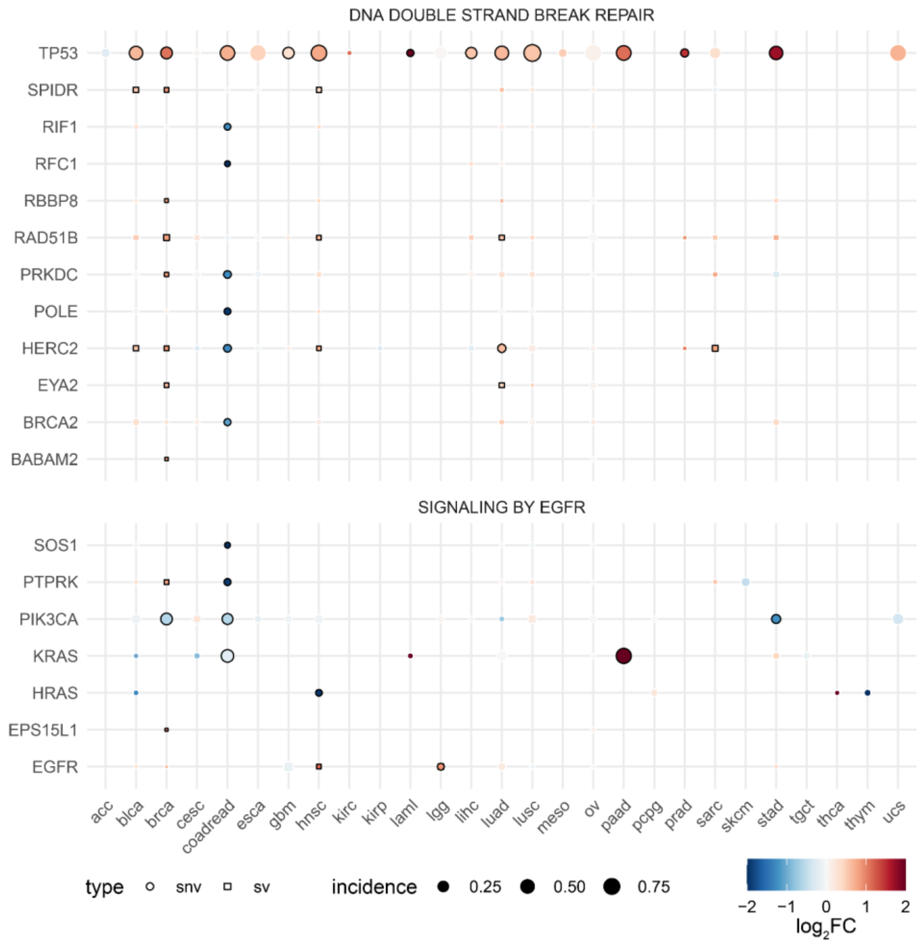


**Figure S18**. Pan-cancer representation of the change in FGA with respect to genomic alterations in genes related to DNA double strand break repair and signaling by EGFR using the Reactome pathway representations. Only genes with a significant association with changes in the FGA in at least one cancer type are shown. Color represents the $log_2$ fold change of the FGA, dot size represents the fraction of samples with a genomic alteration in the corresponding gene, and the dot shape represents the type of genomic alteration affecting the gene. The border of the dots has been colored black when the analysis was significant with an adjusted p-value < 0.05. ACC: Acute Myeloid Leukemia, BLCA: Bladder Urothelial Carcinoma, BRCA: Breast invasive carcinoma, CESC: Cervical squamous cell carcinoma and endocervical adenocarcinoma, CHOL: Cholangiocarcinoma, COADREAD: Colon and Rectum adenocarcinoma, DLBC: Lymphoid Neoplasm Diffuse Large B-cell Lymphoma , ESCA: Esophageal carcinoma, GBM: Glioblastoma multiforme, HNSC: Head and Neck squamous cell carcinoma, KICH: Kidney Chromophobe , KIRC: Kidney renal clear cell carcinoma, KIRP: Kidney renal papillary cell carcinoma, LAML: Acute Myeloid Leukemia, LGG: Brain Lower Grade Glioma, LIHC: Liver hepatocellular carcinoma, LUAD: Lung adenocarcinoma, LUSC: Lung squamous cell carcinoma, MESO: Mesothelioma , OV: Ovarian serous cystadenocarcinoma, PAAD: Pancreatic adenocarcinoma, PCPG: Pheochromocytoma and Paraganglioma, PRAD: Prostate adenocarcinoma, SARC: Sarcoma, SKCM: Skin Cutaneous Melanoma, STAD: Stomach adenocarcinoma, TGCT: Testicular Germ Cell Tumors, THCA: Thyroid carcinoma, THYM: Thymoma, UCEC: Uterine Corpus Endometrial Carcinoma, UCS: Uterine Carcinosarcoma, UVM: Uveal Melanoma. BLCA: Bladder Urothelial Carcinoma, BOCA: Bone Cancer, BTCA: Biliary tract cancer, CLLE: Chronic Lymphocytic Leukemia, CMDI: Chronic Myeloid Disorders, DLBC: Lymphoid Neoplasm Diffuse Large B-cell Lymphoma, EOPC: Early Onset Prostate Cancer, ESAD: Esophageal Adenocarcinoma, GACA: Gastric Cancer, KICH: Kidney Chromophobe, LICA: Liver Cancer, LINC: Liver Cancer, LIRI: Liver Cancer, MALY: Malignant Lymphoma, MELA: Melanoma, ORCA: Oral Cancer OV:

Ovarian serous cystadenocarcinoma, PACA: Pancreatic Cancer, PAEN: Pancreatic Cancer Endocrine neoplasms, PBCA: Pediatric Brain Cancer, RECA: Renal Cancer, UCEC: Uterine Corpus Endometrial Carcinoma.

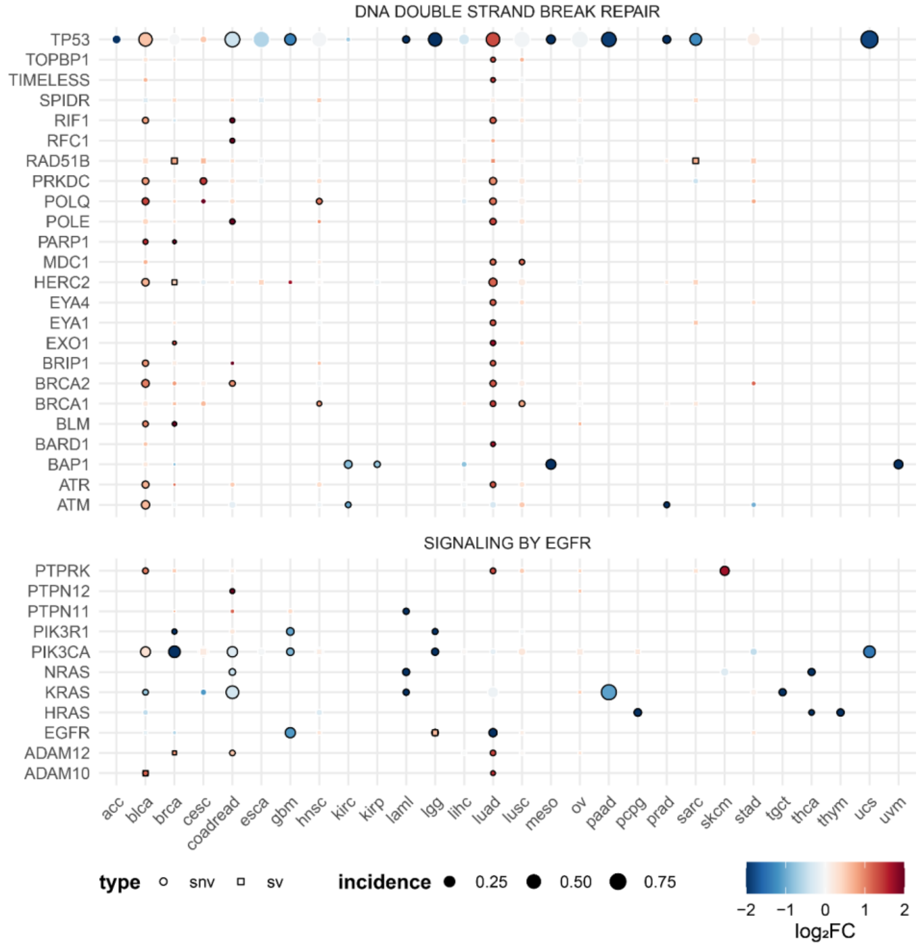


**Figure S19.** Pan-cancer representation of the change in TMB with respect to genomic alterations in genes related to DNA double-strand break repair and signaling by EGFR using the pathway representations of Reactome. Only genes with a significant association with changes in TMB in at least one cancer type are shown. Color represents the $log_2$ fold change of the TMB, dot size represents the fraction of samples with a genomic alteration in the corresponding gene, and the dot shape represents the type of genomic alteration affecting the gene. The border of the dots has been colored black when the analysis was significant with an adjusted p-value < 0.05. ACC: Acute Myeloid Leukemia, BLCA: Bladder Urothelial Carcinoma, BRCA: Breast invasive carcinoma, CESC: Cervical squamous cell carcinoma and endocervical adenocarcinoma, CHOL: Cholangiocarcinoma, COADREAD: Colon and Rectum adenocarcinoma, DLBC: Lymphoid Neoplasm Diffuse Large B-cell Lymphoma , ESCA: Esophageal carcinoma, GBM: Glioblastoma multiforme, HNSC: Head and Neck squamous cell carcinoma, KICH: Kidney Chromophobe , KIRC: Kidney renal clear cell carcinoma, KIRP: Kidney renal papillary cell carcinoma, LAML: Acute Myeloid Leukemia, LGG: Brain Lower Grade Glioma, LIHC: Liver hepatocellular carcinoma, LUAD: Lung adenocarcinoma, LUSC: Lung squamous cell carcinoma, MESO: Mesothelioma , OV: Ovarian serous cystadenocarcinoma, PAAD: Pancreatic adenocarcinoma, PCPG: Pheochromocytoma and Paraganglioma, PRAD: Prostate adenocarcinoma, SARC: Sarcoma, SKCM: Skin Cutaneous Melanoma, STAD: Stomach adenocarcinoma, TGCT: Testicular Germ Cell Tumors, THCA: Thyroid carcinoma, THYM: Thymoma, UCEC: Uterine Corpus Endometrial Carcinoma, UCS: Uterine Carcinosarcoma, UVM: Uveal Melanoma. BLCA: Bladder Urothelial Carcinoma, BOCA: Bone Cancer, BTCA: Biliary tract cancer, CLLE: Chronic Lymphocytic Leukemia, CMDI: Chronic Myeloid Disorders, DLBC: Lymphoid Neoplasm Diffuse Large B-cell Lymphoma, EOPC: Early Onset Prostate Cancer, ESAD: Esophageal Adenocarcinoma, GACA: Gastric Cancer, KICH: Kidney Chromophobe, LICA: Liver Cancer, LINC: Liver Cancer, LIRI: Liver Cancer, MALY: Malignant Lymphoma, MELA: Melanoma, ORCA: Oral Cancer OV:

Ovarian serous cystadenocarcinoma, PACA: Pancreatic Cancer, PAEN: Pancreatic Cancer Endocrine neoplasms, PBCA: Pediatric Brain Cancer, RECA: Renal Cancer, UCEC: Uterine Corpus Endometrial Carcinoma.

**Table S1.** Tumor Break Load (TBL) calculated for three datasets: TCGA, PCAWG and CCLE with the sample ids and tumor type specified.

**Table S2.** Full genomic alteration association analysis results between the genomic instability measures and somatic alterations using TCGA data. The results are reported in three sheets separated on the genomic instability measures TMB, FGA and TBL. The column gene contains the gene names in Hugo gene symbols. Alteration_type indicates the type of somatic alteration from single nucleotide variant (SNV), somatic copy number aberration (SCNA) and structural variant (SV). Cancer type is reported with the TCGA abbreviations. The p-values are reported from a two-sided Mann-Whitney U test with number_cases and number_controls indicating the sample size of the test. P-values are corrected for multiple testing correction using a FDR correction and reported as adjusted p-values. The $\log_2$ foldchange between the mutated and WT groups in terms of genomic instability measure are reported in the column "log2fc". The absolute $\log_2$ foldchange has been used as ranking measure to order the table. Adjusted p-values < 0.05 are deemed significant in this analysis.

**Table S3**. Full genomic alteration association analysis results between the genomic instability measures and somatic alterations using PCAWG data. The results are reported in three sheets separated on the genomic instability measures TMB, FGA and TBL. The column gene contains the gene names in Hugo gene symbols. Alteration_type indicates the type of somatic alteration from single nucleotide variant (SNV), somatic copy number aberration (SCNA) and structural variant (SV). Cancer type is reported with the TCGA abbreviations. The p-values are reported from a two-sided Mann-Whitney U test with number_cases and number_controls indicating the sample size of the test. P-values are corrected for multiple testing correction using a FDR correction and reported as adjusted p-values. The $\log_2$ foldchange between the mutated and WT groups in terms of genomic instability measure are reported in the column "log2fc". The absolute $\log_2$ foldchange has been used as ranking measure to order the table. The ranking measure of genes that are not significant has been set to 0. Adjusted p-values < 0.05 are deemed significant in this analysis.

**Table S4**. Overview of the hazard rate (HR), their 95% confidence interval (95% CI) and the corresponding p-value for high TBL, FGA, or TMB compared to low TBL, FGA or TMB assessed using disease-free survival data from localized (stage I-III) MSS cancer data with known treatment information per cancer type using a multivariate Cox proportional hazards model corrected for age, sex, and tumor stages. Entries with a p-value < 0.05 are highlighted in bold. Hazard rates with an infinite confidence interval are exploded from the table and marked by the"-". Cancer type abbreviations are denoted with the TCGA study abbreviations as provided in TCGA Study Abbreviations | NCI Genomic Data Commons (cancer.gov).

| **cancer type** | **TMB treated DFS** | | **FGA treated DFS** | | **TBL treated DFS** | |
|---|---|---|---|---|---|---|
| | HR (95% CI) | p-value | HR (95% CI) | p-value | HR (95% CI) | p-value |
| BLCA (n = 49) | 1.55 (0.31 - 7.71) | 5.91E-01 | - | 9.98E-01 | 1.69 (0.5 - 5.7) | 3.96E-01 |
| BRCA (n = 739) | **0.54 (0.32 - 0.91)** | **2.02E-02** | **1.84 (1.1 - 3.1)** | **2.11E-02** | **4.24 (1.33 - 13.59)** | **1.49E-02** |
| COADREAD (n = 78) | 5.55 (0.68 - 45.15) | 1.09E-01 | **5.51 (1.34 - 22.66)** | **1.79E-02** | **10.75 (1.21 - 95.59)** | **3.31E-02** |
| ESCA (n = 41) | 1.87 (0.62 - 5.66) | 2.68E-01 | 2.19 (0.59 - 8.06) | 2.39E-01 | - | 9.99E-01 |
| HNSC (n = 61) | 1.3 (0.34 - 5) | 7.02E-01 | **0.29 (0.09 - 0.97)** | **4.43E-02** | 0.18 (0.02 - 1.43) | 1.05E-01 |
| LIHC (n = 36) | 0.97 (0.4 - 2.35) | 9.42E-01 | 4.17 (0.87 - 19.97) | 7.43E-02 | 1.11 (0.49 - 2.53) | 8.03E-01 |
| LUAD (n = 96) | **0.52 (0.27 - 1)** | **4.94E-02** | 0.66 (0.34 - 1.31) | 2.35E-01 | 1.48 (0.72 - 3.06) | 2.88E-01 |
| LUSC (n = 110) | 0.56 (0.19 - 1.63) | 2.85E-01 | 0.29 (0.07 - 1.24) | 9.47E-02 | 0.6 (0.3 - 1.17) | 1.35E-01 |
| PAAD (n = 47) | 1.4 (0.48 - 4.08) | 5.32E-01 | 2.12 (0.61 - 7.35) | 2.38E-01 | **5.13 (1.39 - 18.91)** | **1.41E-02** |
| STAD (n = 96) | 0.67 (0.25 - 1.82) | 4.35E-01 | 0.54 (0.24 - 1.2) | 1.31E-01 | 0.51 (0.22 - 1.21) | 1.27E-01 |
| THCA (n = 193) | **3.4 (1.08 - 10.73)** | **3.64E-02** | **3.19 (1.15 - 8.86)** | **2.60E-02** | 1.26 (0.48 - 3.36) | 6.39E-01 |

**Table S5**. Overview of the hazard rate (HR), their 95% confidence interval (95% CI) and the corresponding p-value for high TBL, FGA or TMB compared to low TBL, FGA or TMB assessed using overall survival data per cancer type using a multivariate Cox proportional hazards model corrected for age, sex and tumor stages. Entries with a p-value < 0.05 are highlighted in bold. Hazard rates with an infinite confidence interval are exploded from the table and marked by the"-". Cancer type abbreviations are denoted with the TCGA study abbreviations as provided in TCGA Study Abbreviations | NCI Genomic Data Commons (cancer.gov).

| | **TMB OS** | | **FGA OS** | | **TBL OS** | |
|---|---|---|---|---|---|---|
| **cancer type** | HR (95% CI) | p-value | HR (95% CI) | p-value | HR (95% CI) | p-value |
| ACC (n = 90) | **5 (2.09 - 11.95)** | **3.00E-04** | **0.31 (0.13 - 0.74)** | **8.35E-03** | **3.51 (1.63 - 7.57)** | **1.37E-03** |
| BLCA (n = 412) | **0.5 (0.37 - 0.7)** | **3.38E-05** | 0.97 (0.71 - 1.32) | 8.32E-01 | **0.58 (0.42 - 0.8)** | **1.05E-03** |
| BRCA (n = 1095) | 1.03 (0.74 - 1.43) | 8.82E-01 | **1.44 (1.01 - 2.06)** | **4.36E-02** | 2.1 (1.44 - 3.05) | 1.08E-04 |
| CHOL (n = 36) | 2.27 (0.77 - 6.66) | 1.35E-01 | **3.51 (1.07 - 11.52)** | **3.86E-02** | - | 9.98E-01 |
| COADREAD (n = 614) | **1.51 (1.02 - 2.22)** | **3.93E-02** | 1.19 (0.81 - 1.74) | 3.75E-01 | **1.46 (1 - 2.11)** | **4.85E-02** |
| ESCA (n = 184) | 1.71 (0.97 - 3.02) | 6.28E-02 | 1.85 (1 - 3.44) | 5.05E-02 | 1.39 (0.82 - 2.37) | 2.23E-01 |
| HNSC (n = 522) | 1.33 (0.99 - 1.8) | 6.08E-02 | 1.1 (0.81 - 1.49) | 5.47E-01 | **1.42 (1.06 - 1.9)** | **2.01E-02** |
| KICH (n = 66) | **9.75 (1.92 - 49.38)** | **5.94E-03** | - | 9.94E-01 | 4.5 (0.83 - 24.38) | 8.11E-02 |
| KIRC (n = 532) | 1.07 (0.78 - 1.48) | 6.67E-01 | 1.17 (0.86 - 1.61) | 3.22E-01 | **1.92 (1.41 - 2.61)** | **3.76E-05** |
| KIRP (n = 290) | **0.2 (0.08 - 0.49)** | **5.31E-04** | **2.13 (1.11 - 4.1)** | **2.34E-02** | **3.05 (1.44 - 6.45)** | **3.63E-03** |
| LIHC (n = 375) | 1.4 (0.95 - 2.07) | 8.96E-02 | **1.6 (1.1 - 2.34)** | **1.51E-02** | 1.12 (0.76 - 1.64) | 5.65E-01 |
| LUAD (n = 518) | 0.92 (0.67 - 1.24) | 5.72E-01 | 1.18 (0.85 - 1.63) | 3.21E-01 | 1.1 (0.81 - 1.49) | 5.52E-01 |
| LUSC (n = 503) | 0.86 (0.64 - 1.16) | 3.22E-01 | **0.73 (0.55 - 0.98)** | **3.54E-02** | 0.79 (0.6 - 1.05) | 1.01E-01 |
| MESO (n = 87) | 1.48 (0.86 - 2.55) | 1.61E-01 | **3.03 (1.8 - 5.12)** | **3.32E-05** | **4.67 (2.47 - 8.83)** | **2.16E-06** |
| PAAD (n = 184) | 1.33 (0.85 - 2.08) | 2.05E-01 | 1.47 (0.87 - 2.49) | 1.53E-01 | **2.01 (1.31 - 3.09)** | **1.42E-03** |
| SKCM (n = 104) | **0.26 (0.1 - 0.65)** | **3.98E-03** | 0.78 (0.34 - 1.79) | 5.54E-01 | **3.35 (1.15 - 9.72)** | **2.64E-02** |
| STAD (n = 442) | **0.64 (0.46 - 0.88)** | **6.87E-03** | 1.33 (0.96 - 1.85) | 8.58E-02 | 1.08 (0.78 - 1.49) | 6.54E-01 |
| THCA (n = 505) | 1.69 (0.54 - 5.27) | 3.64E-01 | 2.24 (0.74 - 6.75) | 1.51E-01 | **5.38 (1.66 - 17.44)** | **5.03E-03** |
| UVM (n = 80) | 1.67 (0.68 - 4.11) | 2.61E-01 | **7.89 (2.26 - 27.5)** | **1.19E-03** | 0.71 (0.29 - 1.74) | 4.48E-01 |
